# Computational functions of precisely balanced neuronal assemblies in an olfactory memory network

**DOI:** 10.1101/2024.04.09.588702

**Authors:** Claire Meissner-Bernard, Bethan Jenkins, Peter Rupprecht, Estelle Arn Bouldoires, Friedemann Zenke, Rainer W. Friedrich, Thomas Frank

## Abstract

Structured connectivity in the brain organizes information by constraining neuronal dynamics. Theoretical models predict that memories are represented by balanced assemblies of excitatory and inhibitory neurons, but the existence and functions of such EI assemblies are difficult to explore. We addressed these issues in telencephalic area Dp of adult zebrafish, the homolog of piriform cortex, using computational modeling, population activity measurements, and optogenetic perturbations. Modeling revealed that precise balance of EI assemblies is important to prevent not only excessive firing rates (“runaway activity”) but also the stochastic occurrence of high pattern correlations (“runaway correlations”). Consistent with model-derived predictions, runaway correlations emerged in Dp when synaptic balance was perturbed by optogenetic manipulations of fast-spiking feedback interneurons. Moreover, runaway correlations were driven by sparse subsets of strongly active neurons, rather than by a general broadening of tuning curves. These results reveal novel computational functions of EI assemblies in an autoassociative olfactory memory network and support the hypothesis that EI assemblies organize information on continuous representational manifolds rather than discrete attractor landscapes.

## INTRODUCTION

Cognition relies on systematic internal representations of knowledge that are thought to be formed by the activity-dependent modification of synaptic connectivity ^1,2^. Representational learning is a main function of autoassociative memory networks, which store relevant information by modifying recurrent synaptic connectivity between specific neuronal assemblies ^1,3^. In classical models of autoassociative memory, assemblies consist of excitatory (E) neurons while the connectivity of inhibitory (I) neurons remains random ^4^. Such assemblies can define stable attractor states and, thus, support the classification of inputs by pattern separation and completion ^5,6^. However, enhanced feedback excitation within assemblies is prone to destabilize networks and generate pathologically high “runaway activity”. Moreover, putative autoassociative brain areas such as hippocampal area CA3 or piriform cortex exhibit activity patterns that are atypical of classical attractor networks such as irregular firing, transient responses to inputs, and high trial-to-trial variability ^7–11^

Biologically realistic firing patterns are generated by networks operating in a regime of inhibition-stabilized synaptic balance (“inhibition-stabilized networks” [ISNs]) ^12–14^. In such networks, individual neurons receive large E and I synaptic inputs that define a membrane potential near spike threshold and generate fluctuation-driven, irregular spike trains. Because small variations in the E/I current ratio cause large firing rate changes, network stability requires co-tuning of E and I synaptic inputs in individual neurons across external stimuli and time, which is referred to as “precise synaptic balance” ^15,16^. E/I co-tuning requires specific higher-order connectivity that can emerge from activity-dependent synaptic plasticity in computational models ^17–19^. Experimentally, E/I co-tuning has been observed in multiple brain areas including sensory cortices ^20–25^ but the underlying network organization remains unclear.

In autoassociative networks, E/I co-tuning may be established by assemblies including both E and I neurons. In such “EI assemblies”, feedback inhibition tracks the activity of E neurons and, thus, curbs runaway excitation without non-specific network-wide suppression of activity ^12,15,18,26^. In ISNs, assemblies do not necessarily establish discrete attractor states but ISNs with structured connectivity may exhibit diverse dynamics including chaotic firing patterns and transient responses ^27–30^. Nonetheless, ISNs with EI assemblies can represent learned inputs and support pattern classification by confining activity to manifolds in activity space ^18^. Generally, recurrent networks with precise synaptic balance can be trained to efficiently perform different computations, conveying a high amount of information per action potential ^15,26,31^. However, computational functions of EI assemblies in ISNs are not fully understood and anatomical evidence is still circumstantial because direct structural analyses of complex network motifs are difficult ^32^.

We addressed these questions in the posterior compartment of telencephalic area Dp (pDp) of adult zebrafish, the homologue of piriform cortex ^33,34^, which is assumed to function as an autoassociative memory network ^35,36^. Dp/piriform cortex are the main targets of the olfactory bulb (OB) and respond to odors with distributed activity that is modified by repeated odor stimulation and learning ^37–41^. Odor-evoked E currents are dominated by recurrent inputs and balanced by inhibition ^20,42–44^. In pDp, voltage clamp recordings directly demonstrated that pDp enters a state of precise synaptic balance during the initial phase of an odor response ^20^.

To explore mechanisms underlying precise synaptic balance in pDp we targeted interneurons contributing to different microcircuits. In piriform cortex, superficial interneurons receive input primarily from the OB and mediate feed-forward inhibition (FFI) whereas deep interneurons receive input from pyramidal neurons and mediate feedback inhibition (FBI)^44–47^. In zebrafish, pDp contains scattered GABAergic interneurons ^37,48^ that have not been characterized in detail. It may be expected that precise synaptic balance depends primarily on interneurons mediating FBI, which tracks population activity, but the identity of these interneurons remains to be determined.

We identified two types of fast-spiking interneurons in pDp that mediated FFI and FBI, respectively, and explored their functions by activity measurements, optogenetic manipulations and network simulations. Using a computational model constrained by data we discovered that the structured connectivity of assemblies can generate high correlations between subsets of input patterns when excitation is not precisely balanced by FBI. These “runaway correlations” can impair pattern classification, occur independently of runaway activity, and depend on stochastic relationships between inputs and assemblies. Consistent with these computational results, optogenetic reduction of inhibition, particularly FBI, generated runaway correlations in pDp by mechanisms consistent with model predictions. These results indicate that precise synaptic balance is important not only to stabilize global activity but also to prevent runaway correlations in recurrent networks with structured connectivity. Moreover, experimental evidence for EI assemblies supports the hypothesis that pDp generates joint representations of odor space and an individual’s experience by confining dynamics to continuous manifolds in activity space ^18^.

## RESULTS

### Genetic targeting of interneuron subtypes

To explore functions of inhibition in a recurrent memory network we genetically targeted two populations of interneurons in pDp using transgenic zebrafish lines (**Fig. 1A**). One line (Tg[SAGFF(LF)212C:Gal4]; abbreviated 212C) was generated in an enhancer/gene trap screen and expressed Gal4 from an insertion near the *ppfia3* locus ^49–51^. The other lines (abbreviated dlx) expressed the Tet trans-activator (itTA) or green fluorescent protein (GFP) under the control of *dlx4/6* promoter/enhancer elements ^52,53^. Previous results showed that dlx elements drive expression in subsets of GABAergic interneurons in the OB of adult zebrafish^54,55^. In the telencephalon, 212C and dlx lines exhibited expression in sparse, largely non-overlapping subsets of neurons (**Fig. 1B**; on average, 6% of 212C^+^ neurons were also dlx^+^, and 7% of dlx^+^ neurons were also 212C^+^; N = 7 fish). Both lines did not show obvious expression in projection neurons and targeted substantially fewer neurons than the gad1b promoter in pDp (Tg[gad1b:GFP], Tg[gad1b:Gal4,UAS:eNpHR3.0YFP]; Frank et al., 2019).

**Figure 1.**
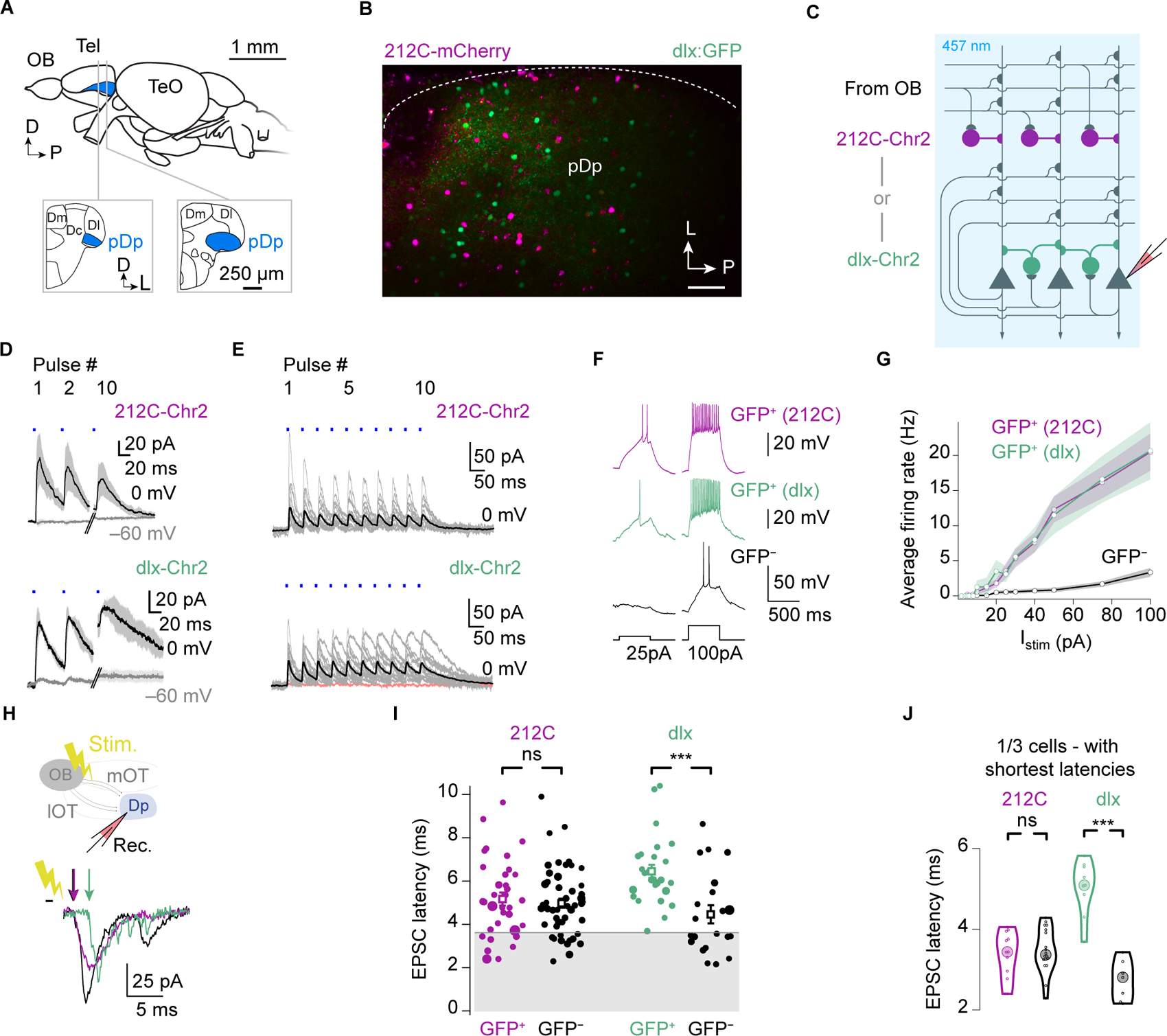
Fast-spiking inhibitory interneurons in pDp. **(A)** Schematic: lateral view of the adult zebrafish brain and two coronal cross sections through the telencephalon (Tel), highlighting the location of pDp. Scale bars represent approximations. OB: olfactory bulb; TeO: optic tectum. Dm, Dc, Dl: medial, central, lateral portions of the dorsal telencephalon, respectively. **(B)** Expression of fluorescent markers in 212C (purple) and dlx neurons (green) in pDp (Tg[SAGFF212C:Gal4,UAS:mCherry,dlx4/6:GFP] fish; maximum intensity projection). Sale bar: 50 µm. **(C)** Schematic: optogenetic stimulation of interneurons and whole-cell patch-clamp recording of evoked IPSCs and EPSCs in putative principal neurons. **(D)** Mean IPSCs (holding potential: 0 mV; ± s.e.m.) and EPSCs (−60 mV) in putative principal neurons in response to 0.5 ms full-field stimulation (457 nm; blue bars) in 212C-Chr2YFP fish (top; n = 4) and dlx-Chr2YFP fish (bottom; n = 2; 10 pulses at 20 Hz; pulses 3 - 9 are not shown). Note absence of optically evoked EPSCs. **(E)** IPSCs (gray: individual neurons; black: average) evoked by trains of blue light pulses (0.5 ms) in putative principal neurons in 212C-Chr2 (n = 11) and dlx-Chr2 fish (n = 16). Only one neuron (red trace) showed no IPSCs. **(F)** Representative firing patterns evoked by current injections (two amplitudes) in interneurons and a putative principal neuron (GFP^−^) in pDp. **(G)** Mean firing rates in the three neuronal populations as a function of injected current. **(H)** Schematic: stimulation of the medial olfactory tract (mOT) and whole-cell recording in pDp. **(I)** Distribution of EPSC latencies in the two interneuron populations (GFP^+^) and the corresponding putative principal neurons (GFP^−^). EPSC latencies of dlx:GFP^+^ neurons, but not 212C-GFP^+^ neurons, were higher than in putative principal neurons (Wilcoxon rank-sum test: dlx:GFP^+^: p = 0.0003, n = 27 vs dlx:GFP^−^, n = 21; 212C-GFP^+^: p = 0.63, n = 34 vs 212C-GFP^−^, n = 53). (**J**) Distribution of 33% shortest EPSC latencies in each neuronal population (subset of data in (H)). Short-latency EPSCs were lacking in dlx:GFP^+^ neurons (Wilcoxon rank-sum test: p = 0.0002; dlx:GFP^+^, n = 9; dlx:GFP^−^, n = 7), but not 212C-GFP^+^ neurons (p = 0.65; 212C-GFP^+^, n = 12; 212C-GFP^−^, n = 18).

To functionally characterize 212C^+^ and dlx^+^ neurons in pDp we crossed drivers to responder lines expressing channelrhodopsin-2-YFP (Chr2YFP) and performed whole-cell voltage clamp recordings in an *ex vivo* preparation of the intact brain and nose (**Fig. 1C**) ^55^. Electrophysiological recordings were performed in the center of pDp and at the boundary between pDp and the *nucleus taeniae* (NT) ^20^. In these regions, 212C-GFP^+^ and dlx:GFP^+^ fibers are abundant but 212C-GFP^+^ and dlx:GFP^+^ somata are sparse. The vast majority of GFP^−^ somata in this region are most likely principal neurons because they do not express the GABAergic marker gad1b-GFP ^37^. We measured E and I postsynaptic currents (EPSCs and IPSCs) in Chr2YFP^-^ neurons held at the reversal potentials of GABAergic and glutamatergic synaptic currents (−60 mV and 0 mV, respectively). Activation of Chr2YFP by trains of full-field blue light pulses (0.5 ms duration; 10 pulses at 20 Hz) evoked prominent IPSCs but no obvious EPSCs in all recorded neurons (**Fig. 1C-E**). Averaged IPSCs showed a weak depression in 212C-Chr2YFP and a weak summation in dlx-Chr2YFP fish (**Fig. S1**). These results show that 212C- and dlx-neurons are inhibitory, presumably GABAergic, interneurons.

When action potentials were evoked in current clamp by depolarizing step currents (500 ms) of increasing amplitude, 212C-GFP^+^ and dlx:GFP^+^ neurons both had lower threshold currents (rheobase) than GFP^−^ neurons (212C-GFP^+^: 34 ± 4 pA, n = 40; dlx:GFP ^+^: 32 ± 4 pA, n = 27; GFP^−^: 122 ± 14 pA, n = 37; Kruskal–Wallis test, n = 104, H = 34.77, p < 10^−8^) and steeper input-output functions (**Fig. 1F, G**). Unlike GFP^−^ neurons, both types of GFP^+^ neurons generated sustained high-frequency trains of action potentials in response to current steps. Moreover, GFP^+^ neurons of both lines had lower action potential thresholds, smaller action potential amplitudes, and shorter action potential durations than GFP^−^ neurons (**Fig. S1**), consistent with previous observations in GABAergic interneurons in Dp ^56^. None of the electrophysiological analyses revealed significant differences between 212C-GFP^+^ and dlx:GFP^+^ neurons, except for a more pronounced after-hyperpolarization following action potentials in dlx:GFP^+^ neurons (p < 0.01; **Fig. S1**). These results indicate that 212C and dlx target distinct types of fast-spiking interneurons in pDp.

Afferent input from mitral cells in the OB may target pDp neurons directly via monosynaptic connections or indirectly via polysynaptic connectivity, which can be distinguished by measurements of synaptic latencies ^57^. We therefore electrically stimulated the medial olfactory tract (mOT) and recorded EPSCs in pDp neurons by targeted voltage clamp recordings ^57^. All neurons showed prominent EPSCs of variable latencies, indicating that synaptic inputs were mono- and polysynaptic. However, short latencies (< 3.8 ms) occurred only in 212C-GFP^+^ and GFP^−^ neurons but not in dlx:GFP^+^ (**Fig. 1I**). When analyzing the shortest 33% of latencies in each dataset, latency distributions were indistinguishable among 212C-GFP^+^, 212C-GFP^−^, and dlx:GFP^−^ neurons but shifted significantly towards longer latencies in dlx:GFP^+^ neurons (p = 0.0002.; **Fig. 1J**). These observations indicate that dlx:GFP^+^ neurons do not receive monosynaptic input from mitral cells. Hence, 212C-GFP^+^ interneurons can provide FFI to other Dp neurons, possibly in combination with FBI, whereas dlx:GFP^+^ interneurons mediate only FBI. 212C-GFP^+^ and dlx:GFP^+^ interneurons are therefore biophysically similar but integrated differently into the synaptic circuitry of Dp.

### Odor-evoked activity of inhibitory interneurons in Dp

We next examined odor responses of 212C and dlx interneurons in Dp. Previous studies indicate that at least some interneurons in piriform cortex and pDp respond selectively to odors^20,58^ but tuning properties of interneurons have not been characterized systematically. We measured odor responses of pDp neurons in 212C-GFP or dlx:GFP fish by 2-photon Ca^2+^ imaging after bolus-loading of the red-fluorescent Ca^2+^ indicator rhod-2 ^41,59^ and inferred action potential firing from the measured fluorescence signals using CASCADE, a pretrained and calibrated artificial network ^60^. This procedure allowed us to directly compare suprathreshold odor responses between 212C-GFP^+^ or dlx:GFP^+^ interneurons and simultaneously recorded GFP^−^ neurons (**Figs. 2A-C**). The stimulus panel comprised twelve structurally diverse odorants including amino acids, bile acids, and nucleotides. Odor stimulation (duration: ∼3 s) evoked robust responses in GFP^−^ neurons and both types of interneurons. The inferred firing rates started to decline before the end of odor stimulation, consistent with previous observations^56,60^

**Figure 2.**
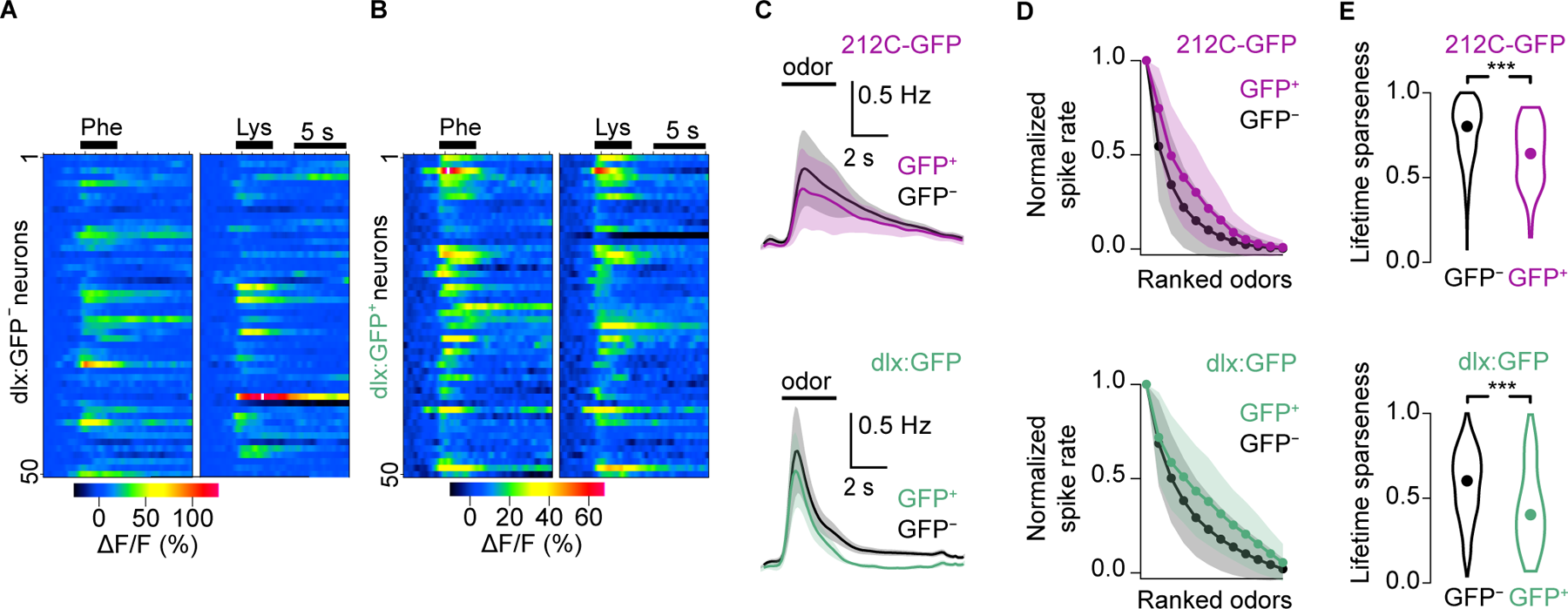
Odor-evoked responses in populations of putative principal cells and interneurons. **(A)** Responses of 50 randomly selected putative principal cells (GFP^−^) in pDp to two amino acid odors (10^−5^ M; average of two trials), measured by 2-photon Ca^2+^ imaging after bolus-loading of rhod-2. **(B)** Odor responses of 50 randomly selected dlx interneurons from the same regions as the GFP^-^ neurons in (A). **(C)** Firing rates inferred from Ca^2+^ signals (see Methods) in interneurons (GFP^+^) and putative principal neurons (GFP^−^), averaged over all trials (n = 2), odors (n = 12) and neurons (212C-GFP: n = 1515 GFP^−^ and n = 50 GFP^+^ from N = 5 fovs; dlx:GFP: n = 1750 GFP^−^ and n = 65 GFP^+^ from N = 12 fovs). Black bar indicates approximate duration of odor stimulation. **(D)** Mean tuning curves of interneurons (GFP^+^) and putative principal cells (GFP^−^) from the same fovs, constructed by rank ordering of odor responses in each neuron. Shading shows s.d. **(E)** Lifetime sparseness of odor responses in interneurons (212C-GFP^+^ and dlx:GFP^+^) and putative principal neurons (GFP^−^; 212C: Wilcoxon rank-sum test: p < 10^−6^; 212C-GFP^+^, n = 50; 212C-GFP^−^, n = 1515; dlx: p < 10^−6^; dlx:GFP^+^, n = 65; dlx:GFP^−^, n = 1750). Lower lifetime sparseness in interneurons indicates broader tuning.

To characterize tuning, we averaged responses of individual neurons over the first 3 s after response onset and sorted them by firing rate to obtain rank-ordered tuning curves. On average, 212C-GFP^+^ and dlx:GFP^+^ neurons were more broadly tuned than GFP^-^ neurons (**Fig. 2D**). Consistent with this observation, odor responses of 212C-GFP^+^ and dlx:GFP^+^ neurons exhibited significantly lower lifetime sparseness than GFP^−^ neurons (212C: p < 0.001; dlx: p < 0.001; **Fig. 2E**) and correlations between odor-evoked activity patterns across 212C-GFP^+^ or dlx:GFP^+^ neurons were significantly higher than correlations across GFP^−^ neurons (**Fig. S2**). However, differences in odor selectivity were modest, and both types of interneurons showed differential responses to odors. Hence, 212C and dlx interneurons exhibited specific responses to different odors although tuning was slightly broader than in principal neurons.

### Effects of inhibitory interneurons on odor-evoked activity in Dp

To examine how interneurons shape odor representations we measured odor responses in transgenic fish expressing the proton pump Archeorhodopsin (212C-ArchTGFP) ^49,61^ or the chloride pump Halorhodopsin (dlx-NpHR3.0YFP) ^62,63^; **Fig. 3**). Odor-evoked Ca^2+^ signals were detected by 2-photon imaging after bolus-loading of Oregon Green 488 BAPTA-1-AM (OGB-1) ^37^. In 50% of the trials, 212C or dlx interneurons were hyperpolarized by targeted illumination of pDp with orange light (594 nm) through an optical fiber for 6.2 s, starting approximately 500 ms prior to odor onset (**Fig. 3A**). In both 212C and dlx fish, photoinhibition of interneurons (PIN) increased odor-evoked population activity, consistent with disinhibition (**Figs. 3B, C**). To quantify changes in firing rate by a measure reflecting feedback gain we normalized the mean firing rate of pDp neurons during PIN (**Fig. 3C**) to the firing rate under control conditions. This “gain index” was calculated for each odor based on the pooled activity of all neurons recorded simultaneously within a given field of view (fov; **Fig. 3D; Methods**). Consistent with disinhibition, the median gain index was >1 in 212C and dlx lines (p < 10^−5^) with a skewed distribution, and larger for PIN of dlx interneurons (212C vs. dlx: p < 10^−4^). We further observed that PIN of 212C or dlx interneurons (PIN_212C_ or PIN_dlx_, respectively) reduced the steepness of rank-ordered tuning curves (**Fig. 3E**), indicating broader tuning. Consistent with these observations, PIN_212C_ and PIN_dlx_ decreased lifetime sparseness (**Figs. 3F**; 212C: p < 10^-16^; dlx: p < 10^-16^; **Fig. 3G**; 212C vs. dlx: p = 0.09) and population sparseness (**Fig. S3**) of odor-evoked activity.

**Figure 3.**
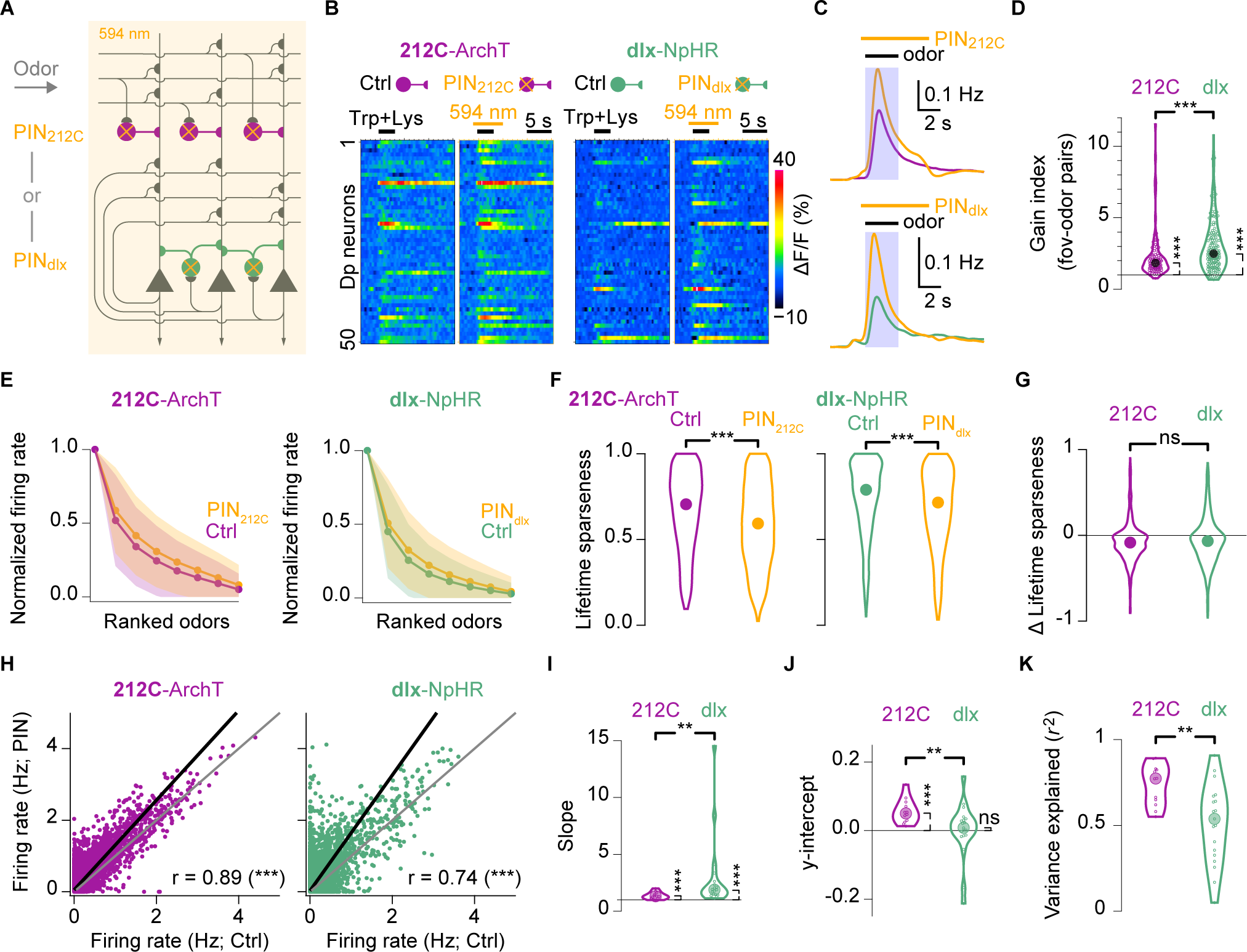
Modulation of odor responses by interneurons in pDp. **(A)** Schematic: photoinhibition (PIN) of 212C or dlx interneurons during odor stimulation and 2-photon Ca^2+^ imaging in 212C-ArchTGFP or dlx-NpHR3YFP fish, respectively. **(B)** Odor-evoked Ca^2+^ signals in 50 randomly selected pDp neurons under control conditions (Ctrl) and during PIN (average of 2 trials). Bars show light exposure and odor stimulation. Left: 212C; right: dlx. **(C)** Inferred firing rates averaged over all neurons, trials and odors under control conditions and during PIN (orange). Bars show light exposure and odor stimulation; shaded area depicts the 3-s time window used for most analyses. Top: 212C; bottom: dlx. **(D)** Gain index (mean activity during vPIN normalized by Ctrl). Each circle represents one fov-odor combination. Gain indices were significantly different from unity (bootstrap test, one-sided; 212C: p < 10^−5^, n = 96; dlx: p < 10^−5^, n = 152) and between 212C and dlx lines (Wilcoxon rank-sum test: p < 10^−4^). **(E)** Rank-ordered odor tuning curves, averaged over all neurons under control conditions and during PIN (orange) in 212C (left) and dlx (right) lines. Shadings show s.d. **(F)** Lifetime sparseness of odor responses under control conditions (left) and during PIN (orange; Wilcoxon signed rank test, two-sided; 212C: n = 1190, p < 10^−15^; dlx: n = 1788, p < 10^−15^) in 212C (left) and dlx (right) lines. **(G)** Distribution of changes in lifetime sparseness (PIN – Ctrl). (**H**) Amplitudes of individual odor responses under control conditions and during PIN. Each data point represents one neuron-odor pair (212C: n = 10,710, N = 12 fovs; dlx: n = 16,092, N = 19 fovs). Black line shows linear fit (total least squares). **(I)** Slopes of linear fits (total least squares) to amplitude data as shown in (H) for each fov (Difference from unity: bootstrap test, one-sided; 212C: p < 10^−5^, N = 12; dlx: p < 10^−5^, N = 19. Difference between lines: Wilcoxon rank-sum test: p = 0.004). **(J)** y-intercepts of the same linear fits (total least squares; Difference from zero: bootstrap test, one-sided; 212C: p < 10^−5^, N = 12; dlx: p = 0.89, N = 19. Difference between lines: Wilcoxon rank-sum test: p = 0.007). **(K)** Variance explained by linear fits. Each datapoint represents one fov (Wilcoxon rank-sum test: p = 0.006).

We next examined whether inhibitory effects were primarily subtractive or divisive. Subtractive inhibition decreases all responses by a constant amount, which sharpens tuning and sparsifies population activity. This form of inhibition has been observed in piriform cortex upon silencing of somatostatin-expressing interneurons ^58^. Divisive inhibition, in contrast, changes the response gain and therefore scales activity patterns without reorganizing their structure. This form of inhibition occurs in dorsal pDp upon non-specific photoinhibition of multiple interneuron types ^37^ and in piriform cortex upon photoinhibition of parvalbumin-expressing interneurons ^58^. Fitting linear functions to odor responses of individual neurons during control conditions and PIN (**Fig. 3H**) yielded slopes >1 for both types of interneurons (212C: m = 1.36 ± 0.09, N = 12 fovs; dlx: m = 2.95 ± 0.75, N = 19 fovs; both p < 10^−5^; **Fig. 3I**). The y-intercept was slightly different from zero for PIN_212C_ but not for PIN_dlx_ (212C: b = 0.06 ± 0.01, p < 10^−5^; dlx: b = 0.00 ± 0.02; p = 0.89; **Fig. 3J**). These observations indicate that dlx and 212C interneurons mediate primarily divisive inhibition. However, linear fits accounted only for a fraction of the variance in the data (212C: 74 ± 4%; dlx: 51 ± 5 %; 212C vs. dlx: p = 0.006; 212C, N = 12 fovs; dlx, N = 19 fovs; **Fig. 3K**). Hence, linear models of uniform divisive and subtractive inhibition cannot fully account for the observed modulation of responses, particularly by PIN_dlx_ (**Fig. 3H, K**), implying that inhibition also has non-uniform effects on odor responses.

### A spiking network model of Dp with feed-forward and feed-back inhibition

To analyze functions of FFI and FBI more systematically we modified a spiking network model of pDp with a single interneuron population ^18^ by introducing two populations of I neurons. The network consisted of 4000 recurrently connected E neurons and a total of 1000 I neurons (**Fig. 4A**), corresponding approximately to the number of neurons in the central region of pDp. E neurons received E input from 1500 afferents (“mitral cells”), consistent with the number of mitral cells in the OB. 500 I neurons received direct E input from mitral cells and mediated FFI while the other 500 I neurons received E input from the E neuron population and mediated FBI (**Fig. 4A**). Both types of interneurons also made I connections within the same population but potential connections between FFI and FBI neurons or additional feed-back connectivity of FFI neurons were not included for simplicity.

**Figure 4.**
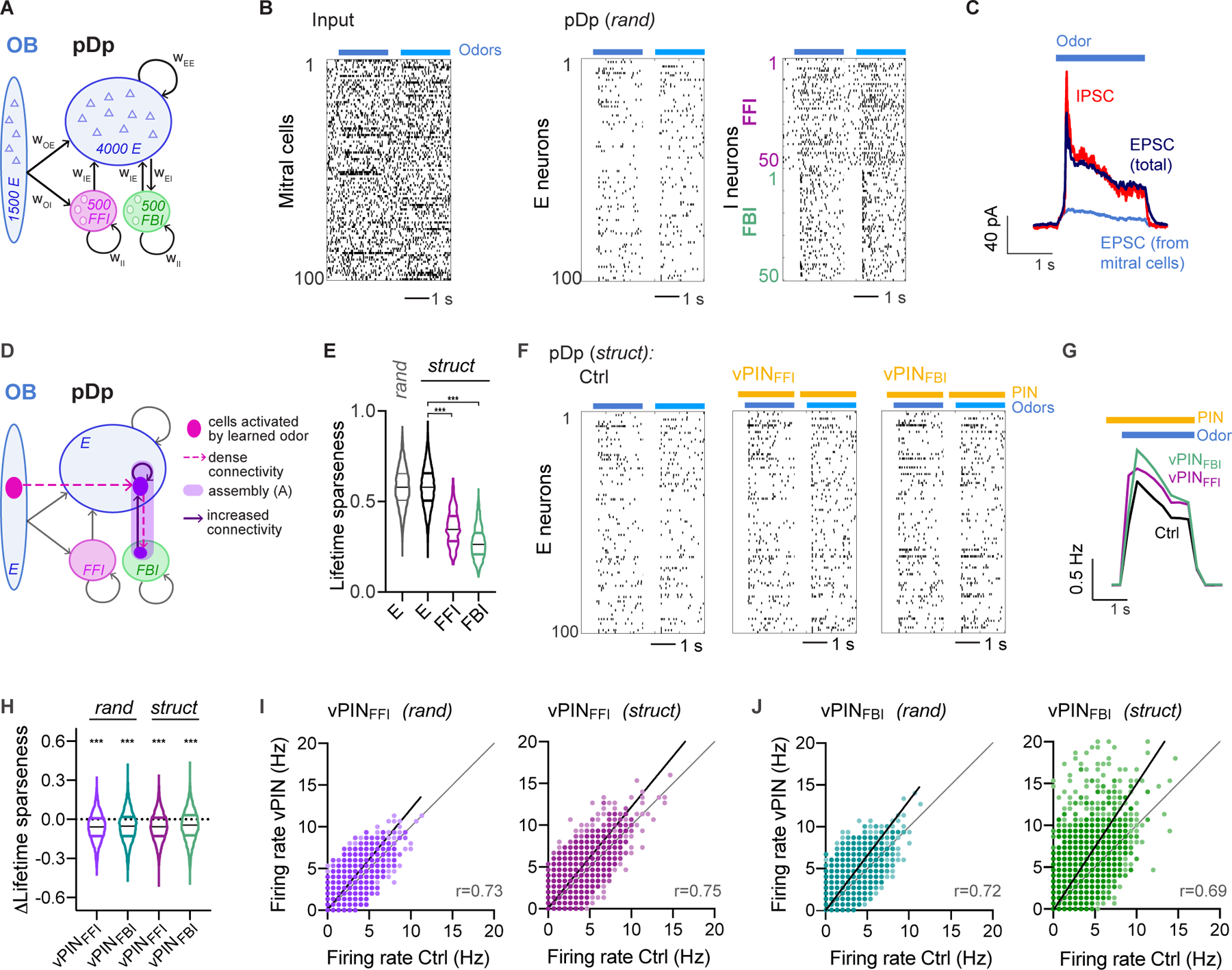
Computational model of pDp. **(A)** Schematic of pDp_sim_. **(B)** Spike raster of randomly selected subsets of 100 mitral cells (input from the OB), 100 E neurons, 50 FFI neurons and 50 FBI neurons. Two different odors (bars) were applied by changing the firing rates of specific subsets of mitral cells for 2 s each. **(C)** EPSCs (black) and IPSCs (red) averaged across all odors and E neurons. Blue trace shows contribution to EPSCs from mitral cells (afferents). **(D)** Illustration of EI assembly in a *struct* network. **(E)** Lifetime sparseness of E neurons in *rand* networks, and E, FFI and FBI neurons in *struct* networks (n = 4000, n = 8000, n = 1000 and n = 1000 from left to right). E neurons were more sharply tuned than FFI (Wilcoxon rank-sum test: p < 0.0001) and FBI neurons (p < 0.0001). **(F)** Spike raster of 100 E neurons in a *struct* network corresponding to the *rand* network in (B): Left: control condition (Ctrl; intact inhibition); center: vPIN_FFI_; right: vPIN_FBi_. **(G)** Mean firing rate averaged over all odors and E neurons under ctrl conditions and during vPIN_FFI_ and vPIN_FBI_. Bars depict odor presentation (blue) and vPIN (yellow). **(H)** PIN-induced change in lifetime sparseness (vPIN – Ctrl; One-sample Wilcoxon signed rank test for difference from zero: p < 0.0001 for all). **(I)** Odor-evoked firing rates of individual neurons during vPIN_FFI_ as a function of their control firing rates in *rand* (left) and *struct* (right) networks (n = 20 odors, 200 neurons, 10 networks) **(J)** Same as (I) for vPIN_FBI_.

Afferent inputs were simulated as Poisson processes with a spontaneous mean rate of 6 Hz. Odor stimuli were modeled as firing rate increases and decreases of 225 and 75 mitral cells, respectively (**Fig. 4B**), mimicking experimental observations in adult zebrafish ^64,65^. Neurons were modeled as conductance-based integrate and fire units with sparse connectivity (connection probabilities ≤10% between all cell types and ≤5% between E neurons). Neuronal parameters were defined based on experimental data when available (membrane time constants; excitatory and inhibitory reversal potentials, spiking thresholds). The remaining neuronal parameters were adjusted to approximate the observed input-output function of E and I neurons in pDp (reset potential, refractory period, firing rate adaptation of E neurons, **Methods**). Connection strengths were then fitted to reproduce experimental observations including a mean odor-evoked firing rate of ∼1 Hz (**Fig. 4B**) ^20,56,59^. Importantly, the network entered a state of synaptic balance during odor stimulation with dominance of recurrent synaptic inputs over afferent inputs (**Fig. 4C**; **Fig. S4**). Very similar activity was generated previously by a simulation of pDp with a single interneuron population ^18^.

To examine the storage of information in synaptic connectivity we simulated two sets of networks: (1) randomly connected networks (*rand*) and (2) structured networks (*struct*) containing EI assemblies with enhanced connectivity. Assemblies were introduced into *rand* networks by increasing connection probabilities among the 60 – 100 E neurons that received most connections from afferents representing a given (“learned”) odor. In addition, the connection probability onto the subset of E neurons was increased from the 10 – 25 FBI neurons that received most connections from these E neurons (**Fig. 4D**, see **Methods** for details). To maintain the number of input connections per neuron, enhanced connectivity within EI assemblies was compensated for by randomly eliminating connections outside assemblies (**Methods**). Including FBI neurons in EI assemblies stabilized the network against runaway activity. This could not be achieved by similar modifications of FFI-to-E neuron connectivity because FFI cannot track the activity of E neurons within the assembly ^12,18^.

We simulated multiple sets of *rand* and *struct* networks starting from different initializations. In each *rand* network, we created EI assemblies representing 20 virtual odors (“learned odors”), resulting in a corresponding *struct* network with 20 memories. Hence, structural odor memories were introduced into the connectivity matrix by enhancing connectivity among small assemblies of E and FBI neurons without modifying FFI. The formation of an assembly involved the modification of only ∼0.15% of the synapses in the corresponding *rand* network. Hence, after creating 20 EI assemblies, ∼3% of synapses were changed, assembly E neurons still received ∼90% of their inputs from neurons outside the assembly, >65% of E neurons remained not affiliated with any assembly, and the majority of connections (∼97%) was shared between *struct* and the corresponding *rand* networks (**Fig. S4B)**.

We then simulated responses of these networks to 20 virtual odors that were different from learned odors, resembling the situation of an experimenter who presents odor stimuli to an animal that has previously formed an unknown set of odor memories (**Fig. S4A**). As observed previously in networks with a single I neuron population ^18^, the mean firing rate, the lifetime sparseness, and other response properties were similar between *rand* and *struct* networks (**Fig. 4E** and **Fig. S4C, D, F**). However, correlated E and I connectivity within assemblies enhanced co-tuning of E and I synaptic currents in individual neurons of *struct* networks (**Fig. S4G**), consistent with experimental observations in pDp ^20^.

We next simulated PIN by reducing FFI or FBI, which enhanced activity of E neurons (**Fig. 4F, G**). Complete elimination of FBI resulted in runaway excitation, which can occur in Dp when inhibition is blocked globally (**Fig. S4H**; ^59^). To mimic experimental observations, we therefore silenced randomly selected subsets of FFI or FBI neurons and adjusted the silenced fraction to approximate PIN-induced increases in population activity (**Fig. 3C; Methods**). Consistent with experimental observations (**Fig. 3H**), this “virtual PIN” (vPIN) revealed primarily divisive inhibition by FFI and FBI neurons. The magnitude of divisive FFI (slope of the linear fit: 1.2) was similar in *rand* and *struct* networks while divisive FBI was more pronounced in *struct* networks (slope: 1.3 in *rand* networks and 1.5 in *struct* networks, **Fig. 4I-J**). As observed in PIN experiments in pDp, linear fits could not fully explain the variance of responses, particularly during vPIN_FBI_ in *struct* networks, implying that inhibition also had non-uniform effects on odor responses (**Fig. 4I-J**). Furthermore, vPIN_FFI_ and vPIN_FBI_ slightly but significantly decreased the lifetime and population sparseness of E neuron responses (**Fig. 4H and Fig. S4I, J**). Simulated networks, particularly those with EI assemblies, therefore reproduced characteristic features of odor-evoked activity in pDp.

### Synaptic balance within EI assemblies prevents runaway correlations

We next examined the transformation of pattern correlations, which quantify the similarity between odor representations. In *rand* networks, correlations between activity patterns across E neurons (“output correlations”) were approximately linearly related to correlations between afferent activity patterns (“input correlations”), consistent with theoretical predictions for random networks ^66,67^. In *struct* networks, output correlations were slightly higher but the linear relationship was largely preserved (**Fig. 5A**). Hence, EI assemblies had only minor effects on global pattern correlations and did not drive pattern separation or completion, as observed in a closely related network simulation with a single interneuron population ^18^.

**Figure 5.**
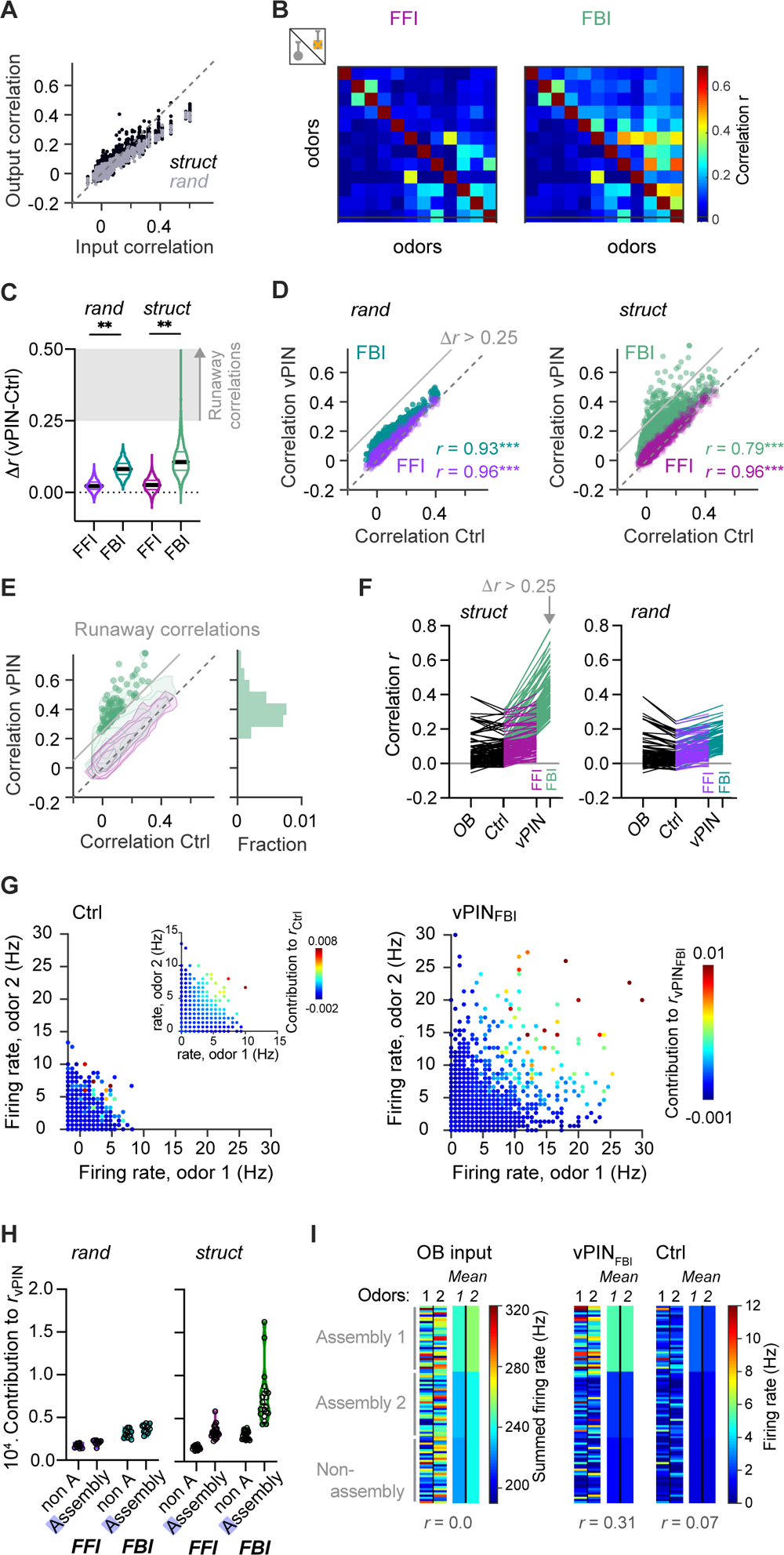
Runaway correlations during imbalanced feedback inhibition in networks with EI assemblies. **(A**) Pearson correlations between odor-evoked activity patterns across E neurons (output correlation) as a function of the correlation between the corresponding afferent activity patterns (input correlation) in *rand* and *struct* networks. **(B)** Example of correlations between activity patterns evoked by 12 odors under control conditions (lower triangles) and during vPIN_FFI_ and vPIN_FBI_ (upper triangles) in one *struct* network. **(C)** Changes in pattern correlations *r* (n = 190 odor pairs) induced by vPIN_FFI_ and vPIN_FBI_ in *rand* and *struct* networks (n = 10 and 20 networks, respectively). vPIN_FBI_ has a stronger effect on pattern correlations than vPIN_FFI_ (Wilcoxon matched-pairs signed rank test; *rand* and *struct*: p < 0.0001). Shaded area depicts 11*r* > 0.25, which was used as an operational criterion to identify runaway correlations. **(D)** Pattern correlations during vPIN_FFI_ and vPIN_FBI_ as a function of correlations under control conditions (Ctrl) in *rand* and *struct* networks. Datapoints above the gray line (11*r* > 0.25) fulfill the operational criterion for runaway correlations. **(E)** Distribution of runway correlations in *struct* networks. Datapoints show runaway correlations (11*r* > 0.25); histogram shows their relative frequency. Contour plots show overall distributions of correlations (same data as in (D); logarithmic contour levels). No runaway correlations occurred during vPIN_FFI_. **(F)** Left: Odor pair-network combinations were selected for occurrence of runaway correlations (11r > 0.25 during vPIN_FBI_) in *struct* networks. For each combination, the corresponding correlations between input patterns, output patterns across E neurons under control conditions (Ctrl) and during vPIN were compared (lines connect datapoints corresponding to the same odor pair-network combinations). Right: correlations for the same odor pair-network combinations in corresponding *rand* networks. High (runaway) correlations were unique to *struct* networks during vPIN_FBI_. **(G)** Contribution of individual neurons to high pattern correlations during vPIN_FBI_. Each datapoint shows the firing rates evoked by two odors in one neuron under control conditions (Ctrl; left) or during vPIN_FBI_ (right) and the neuron’s contribution to the corresponding pairwise pattern correlation during vPIN_FBI_ (color-code). Inset: same as Ctrl but datapoints color-coded by the neuron’s contribution under control conditions. Right: firing rate during vPIN_FBI_. N = 87 odor pairs from 10 networks, 80 randomly selected neurons for each odor pair (Methods). Sparse sets of neurons that responded strongly to both odors made large contributions to pattern correlations and were observed during vPIN_FBI_ but not under control conditions. **(H)**. Contribution of E neurons to all pattern correlations, averaged over neurons that are part of an assembly and neurons outside assemblies (non-A.). **(I)**. Example of activity patterns evoked by two dissimilar odors across E neurons from two assemblies and outside assemblies (left panels: 30 randomly selected neurons per group; right panels: mean activity of each group). Left: input. Assembly 1 received stronger mean input from both odors than assembly 2 and neurons outside assemblies. Right: activity under control conditions (Ctrl) and during vPIN_FBI_. Note increased pattern correlation *r* due to nonlinear amplification of activity in assembly 1 during vPIN_FBI_.

vPIN_FFI_ had minor effects on output correlations in both *rand* and *struct* networks (Δ*r* = *r*_PIN_ - *r*_Ctrl_ = 0.02) while vPIN_FBI_ modestly increased output correlations (**Fig. 5B, C**). Although the mean effect of vPIN_FBI_ was similar in *rand* (Δ*r* = 0.06) and *struct* networks (Δ*r* = 0.08), distributions of Δr differed substantially. In *struct* but not *rand* networks, vPIN_FBI_ strongly increased a subset of output correlations (**Fig. 5D, E**). While Δr never exceeded 0.2 in *rand* networks (n = 1900 odor pair/network combinations), Δ*r* was >0.2 in ∼10 % of the cases and >0.25 in ∼2.5% of cases in *struct* networks. Hence, vPIN_FBI_ resulted in high correlations between a subset of odor representations. Such “runaway correlations”, which occurred in the absence of excessively high “runaway activity”, can impair the storage of independent memories.

To understand how runaway correlations are generated we first focused on pattern correlations that underwent a large increase upon vPIN_FBI_ (Δ*r* > 0.25). In this subset of input/network combinations, the mean pattern correlation during vPIN_FBI_ was *r* = 0.41 ± 0.11; mean ± SD; **Fig. 5F**). Under control conditions, the corresponding pattern correlations were substantially lower (*r* = 0.09 ± 0.08), similar to the corresponding input correlations (*r* = 0.08 ± 0.09), and similar to pattern correlations that were not strongly increased by vPIN_FBI_ (Δ*r* ≤ 0.25, *r* = 0.06 ± 0.08, **Fig. 5F**). Hence, runaway correlations could not be predicted from input correlations or from output correlations when FBI was intact. Runaway correlations are therefore a consequence of structured connectivity when excitation is not precisely balanced by FBI.

We further observed that the main contributions to high pattern correlations in *struct* networks during vPIN_FBI_ came from small subsets of E neurons that responded strongly to both odors in a pair, rather than from weakly responsive, non-selective subpopulations (**Fig. 5G**). These E neurons with high contributions to pattern correlations were dominated by assembly neurons (**Fig. 5H**). Moreover, when odor-evoked input to an assembly was high, activity within the assembly was amplified more than in other assemblies or outside assemblies (**Fig. 5I**). We therefore reasoned that vPIN_FBI_ causes runaway correlations when two odors provide strong input to overlapping sets of assemblies because the reduction of FBI results in a nonlinear amplification of activity within these assemblies (**Fig. S5**).

This hypothesis relies on two basic assumptions: First, inputs should be more strongly amplified within assemblies than outside assemblies because excitatory feedback connectivity is denser within assemblies. Second, the amplification of activity (gain) should increase with the total input to an assembly because the number of neurons that become suprathreshold – and thus contribute to amplification – increases with input strength. Pattern correlations may therefore be enhanced when stimuli activate common sets of assemblies because the amplification of overlapping pattern components will then exceed the mean amplification. Because assemblies are small, the overlap between activity across all neurons is not a strong predictor of the overlap between the activation of assemblies. Nonlinear amplification within assemblies may thus generate high output correlations even when the global input correlation is low. Hence, runaway correlations may occur stochastically depending on the structure of input patterns and network connectivity when excitation within assemblies is not precisely balanced by FBI (**Fig. S5**).

To examine whether this hypothesis can account for runaway correlations we first analyzed firing rates of E neurons within assemblies of *struct* networks as a function of input strength (total firing rate of afferent inputs to the assembly). For comparison, we analyzed firing rates across the same neurons in the corresponding *rand* networks (“pseudo-assemblies”). Firing rates were higher in assemblies than in the corresponding pseudo-assemblies, particularly when input strength was high (**Fig. 6A**). Moreover, vPIN_FBI_ predominantly enhanced strong inputs in assemblies but not in pseudo-assemblies (**Fig. 6A**). These results support the assumptions that the amplification of inputs is higher within assemblies than outside, and that amplification increases nonlinearly as a function of input strength when precise E-I balance is perturbed.

**Figure 6.**
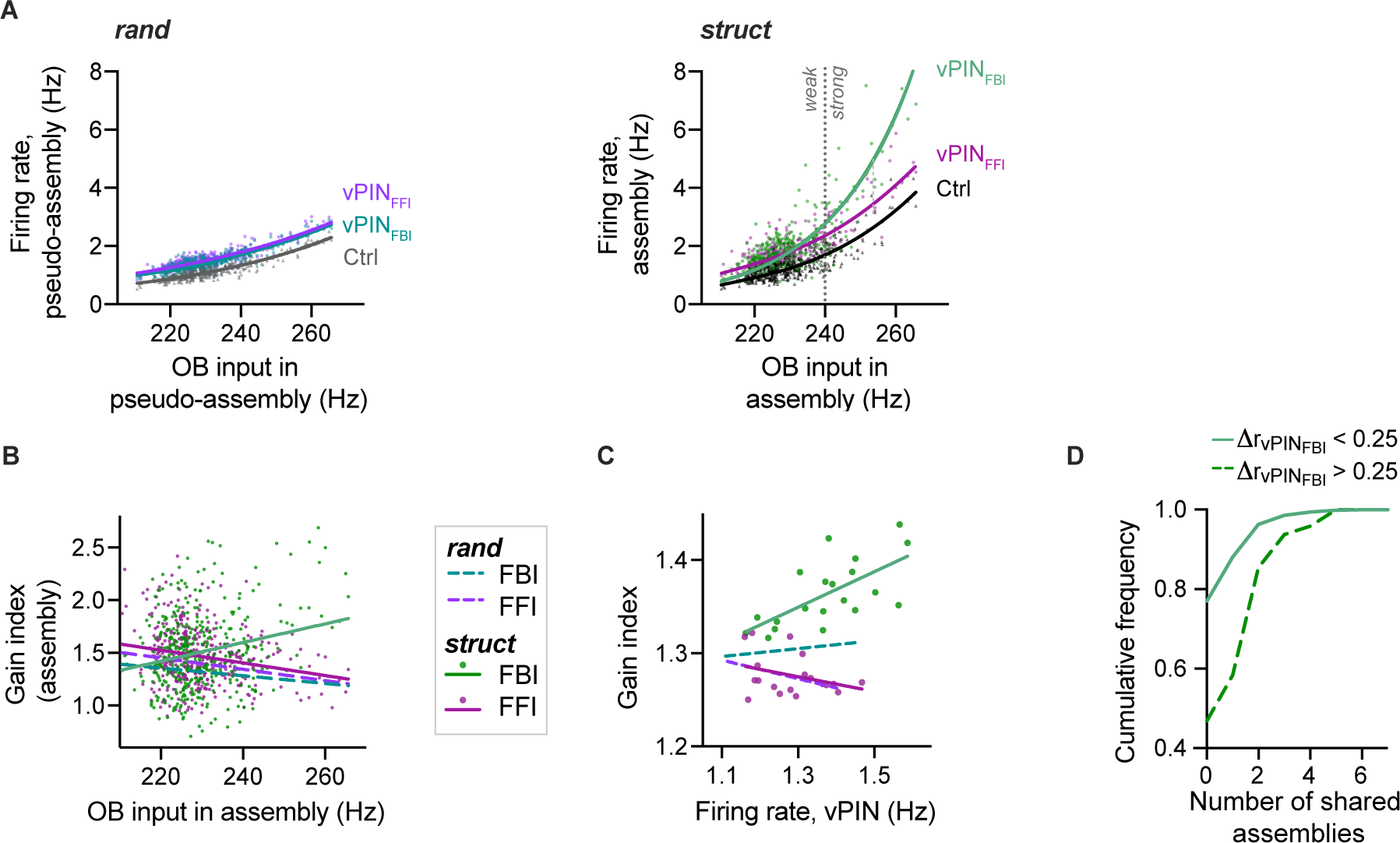
Non-linear amplification in assemblies. **(A)** Mean firing rate of assembly E neurons or the corresponding pseudo-assembly neurons as a function of the mean afferent input to (pseudo-) assemblies (summed firing rates of all connected mitral cells). Data from one *rand* and the corresponding *struct* network with 20 assemblies (mean response to 20 odors). Lines show exponential fits; vertical dotted line shows threshold for “activation” in (D) (240 Hz). **(B)** Gain index (mean activity of assembly during vPIN normalized by Ctrl) as a function of input strength in assemblies (same *rand* and *struct* networks as in (A)). Each datapoint corresponds to one assembly-odor pair in the *struct* network (*rand* not shown for clarity). Lines are linear fits (*rand*: FFI: r = −0.34, p < 0.0001: FBI: r = −0.25, p<0.0001; *struct*: FFI: r = −0.18, p = 0.0003; FBI: r = 0.11, p=0.023). **(C)** Gain index as a function of the mean Dp firing rate during vPIN. Each datapoint corresponds to the gain index in response to one odor averaged over 20 *struct* networks (*rand* not shown for clarity). Lines are linear fits (*rand*: FFI: r = −0.23, p = 0.33; FBI: r = 0.46, p = 0.04; *struct*: FFI: r = −0.22, p = 0.35; FBI: r = 0.72, p = 0.0003). **(D)** Cumulative frequency of shared assembly activation in odor pair-network combinations that exhibited runaway correlations during vPIN_FBI_ (11*r* > 0.25; dashed line) or not (solid line; *struct* networks only). An assembly was defined as “activated” by an odor when the total afferent input to the assembly exceeded 240 Hz (strong input, vertical line in (A)).

We further characterized effects of inhibition by quantifying the gain index (ratio of mean firing rates during vPIN and control conditions). In assemblies of *struct* networks, the gain index for FFI decreased slightly with input strength whereas the gain index increased for FBI (**Fig. 6B**). In the corresponding pseudo-assemblies, in contrast, the gain index of FBI decreased with input strength, as observed for FFI. Similar observations were made also when activity was averaged across all E neurons, rather than within assemblies (**Fig. 6C**). These results confirm that the feedback gain increases with input strength when recurrent excitation is not counterbalanced by specific feedback inhibition.

We next examined whether runaway correlations were associated with the strong activation of overlapping sets of assemblies. An assembly was defined as “strongly activated” when its mean afferent input exceeded a given threshold (**Fig.6A**). We found that vPIN_FBI_ often produced large increases in correlations (Δ*r* > 0.25) when odors shared strongly activated assemblies but rarely otherwise (**Fig. 6D**), supporting the conclusion that runaway correlations depend primarily on the overlap between activated assemblies.

### Functional signatures of EI assemblies in pDp

To examine whether pDp exhibits signatures of EI assemblies we experimentally tested model-derived predictions. We first tested the general hypothesis that FBI counteracts nonlinear recurrent amplification. If so, the gain index during PIN_FBI_ should increase with stimulus strength. To test this prediction, we electrically stimulated the mOT (2 s; 10 Hz) in bulbectomized brain explants (**Fig. 7A; Methods**), measured responses of pDp neurons by 2-photon Ca^2+^ imaging, and transformed Ca^2+^ signals into firing rate estimates using CASCADE^60^. This approach allowed us to measure responses to stimuli of different amplitude under control conditions and during PIN in interleaved trials. As observed during odor stimulation (**Fig. 3B, C**), PIN_212C_ and PIN_dlx_ increased evoked activity (**Fig. 7B-E**). For PIN_212C_, the gain index remained within a narrow range, indicating that responses were scaled by a similar factor, independent of stimulus intensity. The gain index for PIN_dlx_, in contrast, was larger and increased with stimulus strength (**Fig. 7F**), as observed in simulations (**Fig. 6C**). Furthermore, at high stimulus intensities, activity outlasted electrical stimulation (**Figs. 7B-E**), consistent with unbalanced recurrent excitation. This sustained activity was strongly enhanced by PIN_dlx_ (**Fig 7C, D**), resulting in large gain indices at high stimulation intensities (**Fig 7F**), but was nearly unaffected by PIN_212C_ (**Fig. 7B, D**). These observations support the hypothesis that FBI mediated by dlx neurons counteracts nonlinear amplification.

**Figure 7.**
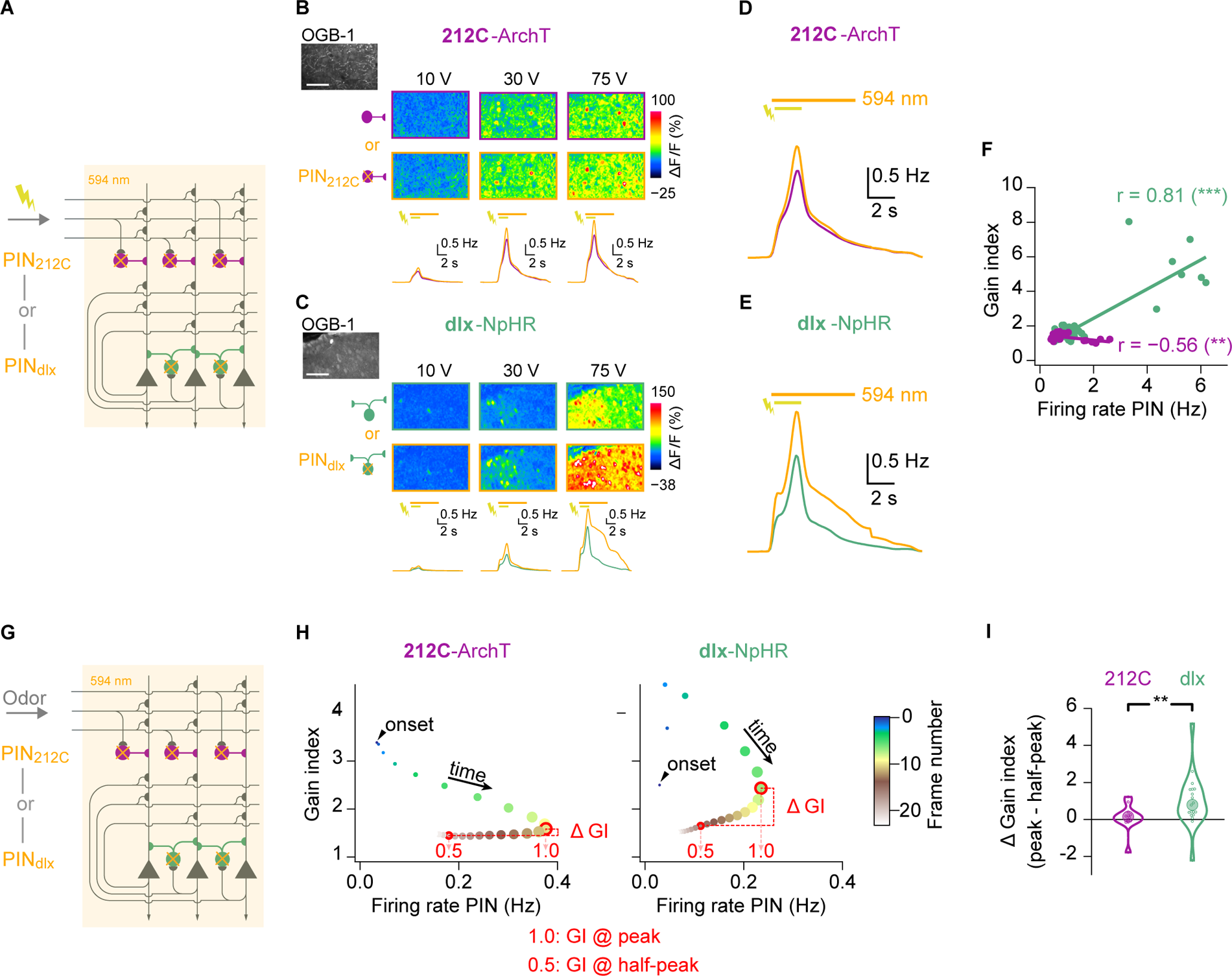
Functional signatures of EI assemblies in pDp: non-linear amplification. **(A)** Schematic: electrical stimulation of medial olfactory tracts (mOT; 20 pulses at 10 Hz; different amplitudes) during simultaneous 2-photon Ca^2+^ imaging and PIN. **(B)** Raw fluorescence of Ca^2+^ indicator (OGB-1) and evoked Ca^2+^ signals (three stimulus amplitudes) under control conditions and during PIN_212C_ (212C-ArchTGFP; representative example; single trials). Bottom row: inferred firing rates averaged over all neurons (n = 903 from N = 12 fovs) and trials (n = 2) for the corresponding stimulus amplitudes under control conditions and during PIN_212C_ (orange). Bars indicate mOT stimulation (2 s) and 594 nm illumination for PIN (6.8 s). Scale bar: 50 µm. **(C)** Same as (B), but for PIN_dlx_ (dlx-eNpHR3.0YFP; n = 2358 neurons from N = 18 fovs). **(D)** Inferred firing rates, averaged over all stimulus amplitudes (n = 6), neurons, and trials under control conditions and during PIN_212C_ (orange). **(E)** Same a (D), but for PIN_dlx_. **(F)** Gain index as a function of inferred firing rate during PIN (212C: r = −0.56, N = 24 fov-stimulus pairs, p = 0.004; dlx: r = 0.83, N = 33 fov-stimulus pairs, p < 10^−8^). **(G)** Schematic: Ca^2+^ imaging of odor responses during PIN. **(H)** Gain index during odor responses as a function of time. The difference in gain index (11 GI) was calculated between the two time points corresponding to the peak of response (“1.0”; open red circle) and the decay to 50% (“0.5”; closed red circle) for each fov-odor pair. Arrowhead indicates onset of odor response. Time is color-coded; marker size is proportional to the inferred firing rate during PIN (x axis). (**I**) Change in gain index during odor responses, calculated as in (G), in different fovs (Wilcoxon rank-sum test: p = 0.009; 212C line: N = 12; dlx line: N = 19).

We next examined signatures of non-linear FBI in responses to odors (**Fig. 7G**). During an odor response, pDp neurons receive synaptic inputs mostly from other E neurons in pDp and potentially other telencephalic areas ^20^. As activity within pDp varies over time (**Figs. 2, 3**), the gain index for FBI should co-vary with this activity if amplification and FBI vary with input strength. To address this prediction, we focused on the time window between the response peak and its decay to 50%, which typically occurred within approximately 2 s (**Fig. 7H**; the initial response transient was not analyzed because activity changed rapidly ^56,59^). While the gain index for PIN_212C_ remained almost constant during this time, the gain index for PIN_dlx_ decreased together with the firing rate (**Fig. 7H**). Consequently, the change in gain index was significantly larger for PIN_dlx_ than for PIN_212C_ (**Fig. 7I**; p < 0.01). These observations further support the conclusion that 212C neurons scale activity by a constant factor whereas dlx neurons balance nonlinear, activity-dependent amplification.

To directly examine whether unbalanced activity generates runaway correlations, we compared correlations between activity patterns evoked by eight different odors under control conditions and during PIN (**Fig. 8A**). The mean pattern correlation was not affected by PIN_212C_ (Δ*r*: 0.04 ± 0.04, mean ± s.e.m., p = 0.21, bootstrap test; **Fig. 8B**) but significantly increased by PIN_dlx_ (Δ*r*: 0.12 ± 0.04, mean ± s.e.m., p < 0.001, bootstrap test; **Fig. 8B**). Large increases in correlations occurred between subsets of activity patterns upon PIN_dlx_ but rarely upon PIN_212C_ (**Fig. 8E, F).** As a consequence, effects of PIN_212C_ and PIN_dlx_ were uncorrelated (**Fig. 8C**), despite a high similarity between correlation matrices under control conditions (**Fig. 8D**). As observed in simulations during vPIN_FBI_, high pattern correlations were driven by small subsets of neurons that responded with high firing rates to both odors during PIN_dlx_ but not under control conditions (**Fig. 8H**). During PIN_212C_, in contrast, high contributions of individual neurons to pattern correlations were rare and not associated with high firing rates (**Fig. 8G**). We therefore conclude that perturbations of FBI can generate runaway correlations in pDp due to changes in the activity of small subsets of neurons, consistent with assembly-driven runaway correlations observed in the computational model.

**Figure 8.**
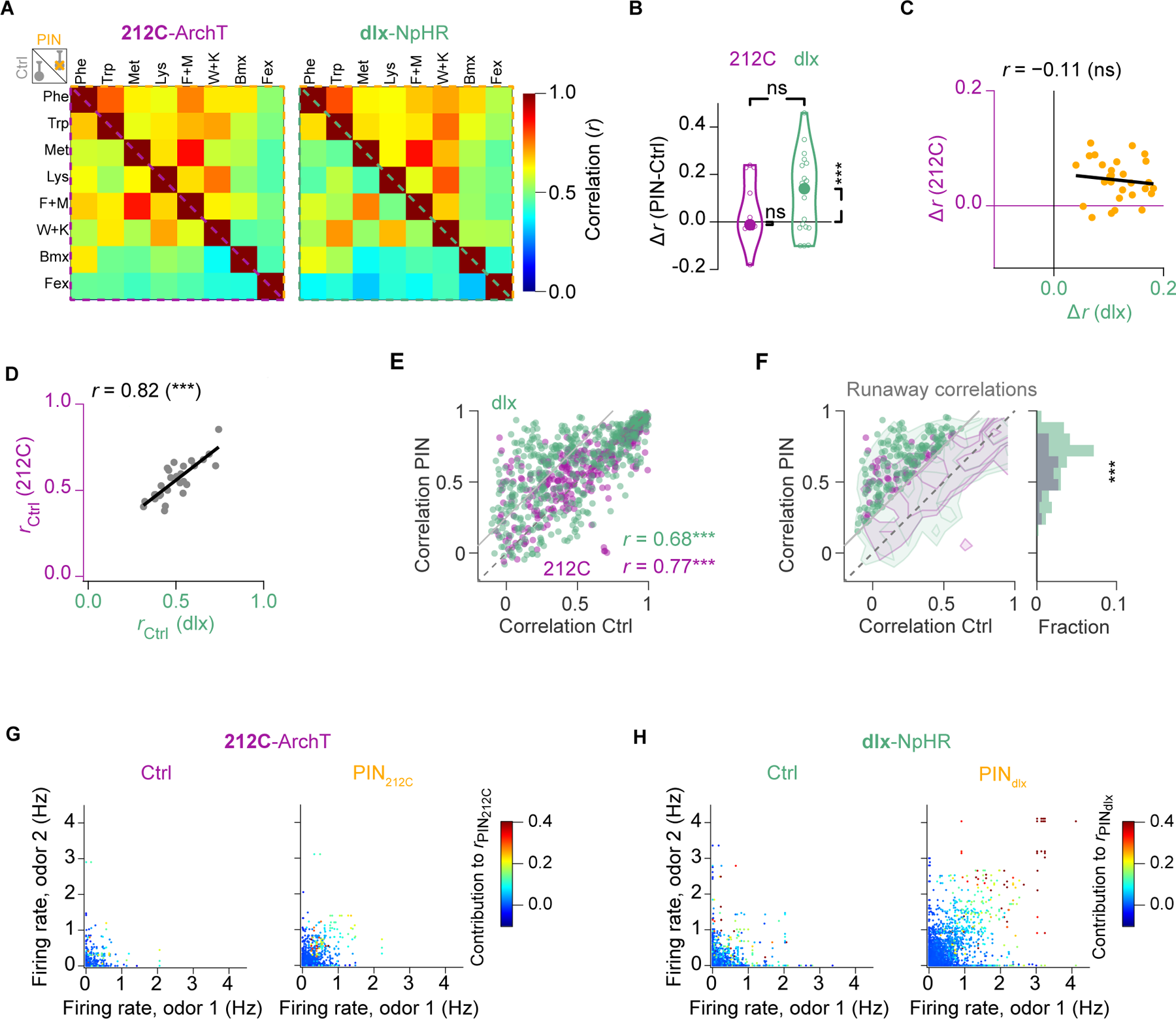
Functional signatures of EI assemblies in pDp: population activity patterns and runaway correlations. **(A)** Average Pearson correlation (*r*) between odor-evoked activity patterns under control conditions (Ctrl, below diagonal) and during PIN (above diagonal; 212C: N = 12 fovs; dlx: N = 19 fovs). **(B)** Mean PIN-induced changes in pattern correlations (Δ*r* = *r*_PIN_ − *r*_Ctrl_; bootstrap test, one-sided; 212C: p = 0.21, N = 12; dlx: p = 0.0005, N = 19; 212C vs dlx, Wilcoxon rank-sum test: p = 0.33). **(C)** Relationship between changes in correlations evoked by the same odors (n = 28 odor pairs) during PIN_212C_ and PIN_dlx_. **(D)** Relation between the mean pattern correlation (n = 28 odor pairs) in 212C and dlx lines under control conditions. **(E)** Pattern correlations during PIN_212C_ and PIN_dlx_ as a function of correlations under control conditions. Datapoints above the gray line (11*r* > 0.25) fulfill the operational criterion for runaway correlations. **(F)** Distribution of runway correlations. Datapoints show runaway correlations (11*r* > 0.25); histogram shows their relative frequency. Contour plots show overall distributions of correlations (same data as in **(E)**; logarithmic contour levels). Frequency of runaway correlations was significantly higher during PIN_dlx_ (p = 10^−9^; Ξ^2^-test). **(G)** Contribution of individual neurons to high pattern correlations. Each datapoint shows the firing rates evoked by two odors in one neuron under control conditions (Ctrl; left) and during PIN_212C_ (right). The neuron’s contribution to the corresponding pairwise pattern correlation during PIN_212C_ is color-coded. **(H)** Same as (G) for PIN_dlx_. Note sparse sets of neurons that responded strongly to both odors and made large contributions to pattern correlations during PIN_dlx_ but not under control conditions.

## DISCUSSION

EI assemblies can establish precise synaptic balance in ISNs, which is thought to stabilize recurrent memory networks against runaway activity ^12,16,18^. However, direct evidence for EI assemblies in biological networks and a comprehensive understanding of their computational functions is lacking. Using computational modeling we found that precise E/I balance is important not only for network stability but also to prevent *runaway correlations* between subsets of inputs. These *runaway correlations* are high pattern correlations driven by sparse subsets of neurons whose firing rates are modest under control conditions but strongly increased when E/I balance is perturbed. To explore functions of EI assemblies in pDp we photoinhibited different types of fast-spiking interneurons and found signatures of EI assemblies primarily during manipulations of FBI, consistent with model-derived predictions. These results show that connectivity motifs generating precise synaptic balance are critical for efficient memory storage in recurrent circuits, and that balanced EI assemblies are likely to mediate memory-related computations in pDp.

### Functional characterization of interneurons in pDp

We characterized two distinct populations of fast-spiking I interneurons with similar biophysical properties, dlx and 212C, that mediate FBI and FFI (possibly in combination with FBI), respectively. IPSCs generated by 212C and dlx neurons exhibited depression and summation, respectively, similar to short-term synaptic plasticity of FFI and FBI interneurons in piriform cortex ^45,46^. During prolonged odor responses, the relative weight of inhibition may therefore shift from FFI to FBI. However, 212C and dlx neurons presumably represent only a subset of all interneurons in pDp.

Optogenetic hyperpolarization of 212C and dlx neurons caused disinhibition without triggering runaway excitation, which allowed us to dissect computational functions of defined interneurons in an ISN. Both interneuron types mediated primarily divisive inhibition, as observed for fast-spiking parvalbumin neurons in piriform cortex ^58^. Divisive inhibition is well-suited to scale and globally stabilize population activity in recurrent networks ^68^. In rodents, FBI has also been proposed to contribute to concentration-invariant odor identity coding by curtailing long-latency afferent input during a sniff cycle ^69–71^. It remains to be determined whether FBI has similar functions in zebrafish given that the kinetics of odor responses is slower and not modulated by sniffing.

While linear models of inhibition could account for most effects of PIN_212C_, PIN_dlx_ had additional neuron- and stimulus-specific effects on odor responses. Consistent with this finding, correlations between odor-evoked activity patterns were modified substantially by PIN_dlx_ but not by PIN_212C_, even though both manipulations increased mean firing by similar amounts. Dlx neurons therefore have non-uniform effects on population activity that suppressed runaway correlations, indicating that they contribute to the specificity of odor representations in pDp.

### Signatures of structured connectivity in inhibition-stabilized networks

Important functions of inhibition include the stabilization of recurrent networks against runaway excitation, the modulation of tuning curves, and temporal patterning of activity ^72^. Our results indicate that a further function of inhibition in memory networks is the suppression of runaway correlations, which can emerge from structured connectivity during learning.

This insight was obtained using an ISN model constrained by data from pDp ^18,20,56,59^. When recurrent excitation was not precisely balanced by FBI, assemblies strongly increased subsets of pattern correlations without generating runaway activity, implying that runaway correlations were not caused by global E/I imbalance. Rather, high correlations were driven by sparse and strong responses of E neurons within specific assemblies (**Fig. S5**). Hence, runaway correlations could not be predicted from global pattern correlations but were a direct consequence of structured connectivity, which is established during learning. In our computational model, the nonlinearity in the amplification of activity within assemblies depended entirely on the threshold-nonlinearity of neuronal input-output functions (action potential generation). In piriform cortex and other brain areas, E neurons contain additional nonlinearities ^73,74^ that are likely to further enhance nonlinear amplification within assemblies. While experience-dependent changes in correlations due to the formation of structured connectivity could, in principle, have different functions ^75^, high correlations generally impair autoassociative memory by limiting discriminability and storage capacity. The suppression of runaway correlations may thus be important, and possibly critical, for the function of recurrent memory networks. It may therefore be expected that runaway correlations are rarely observed in biological memory networks without perturbations of E/I balance. Nonetheless, runaway correlations may occur naturally under certain conditions of imprecise E/I balance, for example during learning, in response to neuromodulatory inputs, or when the computational function of a network is supported by high correlations.

Coordinated E-I connectivity underlying precise synaptic balance may be established by different motifs including EI assemblies, which can be generated by biologically plausible learning rules ^19,76^. However, experimental evidence for EI assemblies remains indirect. In primary visual cortex, for example, interactions between E and I neurons depend on orientation tuning ^77–80^ but the statistical knowledge obtained by sparse sampling of neurons is insufficient to resolve higher-order connectivity motifs in large networks ^81,82^. We therefore examined diagnostic features of EI assemblies in population activity by manipulations of inhibitory interneurons. Runaway correlations were observed primarily during PIN_dlx_, consistent with a suppression of runaway correlations by FBI under control conditions. PIN_dlx_-induced runaway correlations were driven by strong responses of sparse neuronal subsets rather than by a general broadening of response selectivity, consistent with the mechanism underlying runaway correlations in simulations. While PIN_212C_ also increased a subset of correlations, possibly due to a feedback component of 212C neurons, effects were smaller and not driven by sparse and strong responses. These results support the hypothesis that precise synaptic balance in pDp ^20^ is established, at least in part, by EI assemblies.

In the OB, principal neurons (mitral cells) receive E input predominantly from sensory afferents while recurrent E connections are very rare ^83,84^. As revealed by a combination of activity measurements and dense circuit reconstructions (“dynamical connectomics”), mitral cells receiving co-tuned sensory input preferentially inhibit each other via reciprocal connectivity with common interneurons, thereby attenuating correlated activity ^83^. This mechanism of pattern decorrelation in the olfactory bulb is similar to the suppression of runaway correlations in EI assemblies although correlated activity of mitral cells is generated by common input rather than recurrent excitation. Pattern decorrelation by specific FBI of correlated activity may thus be a common computational motif in different types of networks.

### Computational functions of EI assemblies in pDp

Piriform cortex and pDp are thought to generate experience-dependent representations of olfactory objects, environments or tasks by autoassociative mechanisms ^35,36,85^. Unlike classical attractor models, piriform cortex and Dp generate transient, variable and irregular firing patterns ^41,56,60,86–88^. These observations were reproduced by a recent ISN model of pDp^18^. This model indicates that EI assemblies project odor representations of learned inputs into activity subspaces by locally confining activity onto manifolds, resulting in continuous representations that nonetheless enhanced classification of learned inputs by assembly neurons. Hence, EI assemblies may cause geometric modifications of coding space that support continuous computations such as navigation or measurements of relevant distances, possibly in the context of cognitive maps.

ISN models generating continuous representational manifolds assume that experience drives the formation of EI assemblies. Our results provide experimental support for these models by providing indirect evidence for EI assemblies in pDp. Representational manifolds combine a sensory map of olfactory stimulus space – transmitted from the olfactory bulb – with an individual’s experience, suggesting that pDp generates representations of odor space with individualized geometries. This hypothesis predicts that learning geometrically modifies (“distorts”) a pre-existing and continuous map of odor space, rather than establishing discrete representations of specific odors. Consistent with this hypothesis, repeated odor stimulation gradually modified odor-evoked activity patterns, and representations of novel and learned odors were not categorically different in pDp ^37,41^. Gradual rather than categorical modifications of odor representations were also observed in piriform cortex as a consequence of passive odor presentation or active learning ^39,85,89^. Continuous representational manifolds may support fast classification and interpretation of inputs, complementary to integrative functions of continuous attractor networks. Future studies may take advantage of the small size of pDp to directly analyze the underlying network structure by combining activity measurements with dense reconstructions of neuronal wiring diagrams ^32^.

## METHODS

### Animals and transgenic lines

Experiments were performed using adult (>3 months) zebrafish (*Danio rerio*) of both sexes. Fish were raised and kept under standard laboratory conditions (26 – 28 °C; 13/11 or 14/10 light/dark cycle). All animals are sacrificed prior to the removal of organs in accordance with the Veterinary Department of the Canton Basel-Stadt (Switzerland) or the European Commission Recommendations for the euthanasia of experimental animals (Part 1 and Part 2). Experiments involved the following transgenic lines:

| **Short name** | **Systematic name** | **References** |
| --- | --- | --- |
| 212C-mCherry | *Tg(SAGFF(LF)212C:Gal4, UAS:mCherry)* | <sup>49</sup>, *this study* |
| 212C-GFP | *Tg(SAGFF(LF)212C:Gal4, UAS:GFP)* | <sup>49,51</sup> |
| dlx:GFP | *Tg(nccr.i56i-i56ii-dlx6a:GFP)* | <sup>52</sup> |
| vglut2a:DsRed | *TgBAC(slc17a6b:LOXP-DsRed-LOXP-GFP)* | <sup>90</sup> |
| 212C-Chr2YFP | *Tg(SAGFF(LF)212C:Gal4, 5xUAS:wtChr2-EYFP)* | <sup>49</sup>, *this study* |
| 212C-ArchTGFP | *Tg(SAGFF(LF)212C:Gal4, 5xUAS:Htp.ArchT-GFP,myl7:EGFP)<sub>fmi13</sub>* | <sup>49</sup> |
| dlx-Chr2YFP | *Tg(dlx5a-dlx6a:ITETA;PTET:Cr.Cop4-YFP)<sub>fmi9</sub>* | <sup>53</sup> |
| dlx-eNpHR3.0YFP | *Tg(5xUAS:ITETA,PTET:eNpHR3-YFP,myl7:EGFP)<sub>fmi14</sub>* | <sup>63</sup> |

The UAS:Chr2YFP expression construct was generated using the Tol2Kit ^91^, which involved a multisite recombination reaction (Invitrogen Multisite Gateway manual v.D, 2007) between p5E–UAS (5×UAS and E1b minimal promoter; Distel et al., 2009), pME–wtChr2YFP; ^92^) and p3E–polyA as entry vectors, and pDestTol2CG2 as destination vector ^91^. Stable transgenic founder lines (UAS:Chr2YFP, UAS:mCherry) were generated using standard procedures ^93^

### Experimental preparation

Electrophysiological measurements, Ca^2+^ imaging, and imaging of fluorescent reporter expression were performed in an ex-vivo preparation of the entire zebrafish brain and nose ^55^. Briefly, adult zebrafish were either cold-anesthetized or, for some experiments to characterize expression patterns, anesthetized by immersion in MS-222 and decapitated. The forebrain was exposed ventrally after removing the eyes, jaws and palate. The preparation was placed in a custom-made flow-chamber, continuously superfused with teleost artificial cerebrospinal fluid (ACSF) and slowly warmed up to room temperature. ACSF contained (in mM): 124 NaCl, 2 KCl, 1.25 KH_2_PO_4_, 1.6 MgSO_4_, 22 D-(+)-Glucose, 2 CaCl_2_, 24 NaHCO_3_, pH 7.2. Chemicals were obtained from Sigma-Aldrich. In some experiments (see below) both olfactory bulbs were surgically removed (“bilateral bulbectomy”) while the rest of the telencephalon remained intact.

### Electrophysiology, optogenetics and olfactory tract stimulation

Voltage clamp and current clamp recordings were performed using patch pipettes that were pulled from 1-mm borosilicate glass capillaries (Hilgenberg) with a resistance of 5–8 MΩ, a Multiclamp 700B amplifier (Molecular Devices) and Ephus software ^94^. Pipettes were filled with an intracellular solution that contained (in mM): 129 K-gluconate, 10 HEPES (free acid), 0.1 EGTA, 4 Na_2_-ATP, 10 Na_2_-phosphocreatine, 0.3 Na-GTP, 5 L-glutathione and 13.1 KOH (pH 7.2, 305 mOsm; all Merck / Sigma). Neurons were targeted by a combination of contrast-enhanced transmitted-light optics and multiphoton fluorescence using the shadow-patching technique with 50 - 100 µM Alexa Fluor 594 or Alexa Fluor 488 (Thermo Fisher Scientific) in the internal solution. Prior to recordings, we usually removed the dura mater over pDp. Before making a seal, neurons were approached with low pressure (approximately 20 mbar). Measurements were not corrected for the liquid junction potential and signals were digitized at 10 kHz after low-pass filtering. All recordings were performed in pDp ^20^. In current clamp recordings (**Fig. 1F, G**; **Fig. S1B-F**), we evoked action potentials by current injection at different amplitudes in 212C-GFP (N = 12 fish) or dlx:GFP fish (N = 7). For the analysis of firing rates as a function of input current, five trials were averaged per cell and amplitude (**Fig. 1F, G**). Detailed analyses of action potentials (**Fig. S1B-F**) were restricted to the first action potential at rheobase.

Optical stimulation of Chr2YFP (**Fig. 1C, D; Fig. S1F, G**) was performed as described^55^ using a digital micromirror device (DMD) that was optically coupled into a multiphoton microscope and illuminated with a blue laser (457 nm, 500 mW unattenuated laser output). “Full-field” optical stimulation (600 x 600 pixels) covered the majority of pDp in its anterior-posterior and medio-lateral extent, and possibly small parts of adjacent areas. Ten light pulses of 0.5 ms duration were delivered at 20 Hz. Inhibitory (IPSCs) and excitatory postsynaptic currents (EPSCs) were recorded from Chr2-negative, vglut2a-positive (excitatory) neurons in pDp using whole-cell patch-clamp recordings at holding potentials of 0 mV or –60 mV, respectively (averages over five trials).

To measure EPSC latencies (**Fig. 1H-J**) we electrically stimulated the medial olfactory tract (mOT) as described ^57^ with minor modifications of previous procedures. We stimulated the mOT unilaterally about 100 µm posterior from the border of the OB using glass pipettes with a tip diameter of ∼10 – 20 µm. Pipettes were filled with 1 M NaCl. Stimulus amplitude (usually ∼−30 V) was adjusted to evoke responses of intermediate amplitude and EPSCs were recorded at –70 mV. Ten stimuli (0.5 ms) were applied at 20 Hz, resulting in ten post-stimulus segments for EPSC analysis. At least five trials were recorded in each neuron, resulting in at least 50 segments per neuron. Latencies of the first EPSC in individual segments were analyzed in a semi-automatic fashion using custom scripts written in IgorPro (Wavemetrics). The procedure first used a published event detection algorithm ^95^ followed by visual inspection and, if necessary, manual correction of the EPSC onset. Importantly, all latency analyses were performed blind with regard to the recorded cell type. To avoid spurious effects of spontaneously occurring EPSCs we averaged latency values for each segment across trials, and report the minimum of these segment averages for each neuron.

### Loading of the Ca^2+^ indicator, odor application, and tract stimulation

Rhod2-AM (**Fig. 2** and **Fig. S2**) or Oregon Green 488 BAPTA-1-AM (OGB-1; other figures; all ThermoFisher Scientific) were bolus-loaded as described ^55^ with minor modifications. 50 μg of AM dye was dissolved in 30 μL of DMSO/Pluronic F-127 (80/20; ThermoFisher Scientific) and stored in 4 μL aliquots at −20°C. Prior to each experiment, an aliquot was diluted 1:5 in ACSF and loaded into a glass pipette with a tip diameter of approximately 5 μm. Pressure injections were targeted to the lateral telencephalon, posterior to the prominent furrow and blood vessel and within or slightly dorsal to Dp. One or a few injections were made up to 100 µm dorsal to the subsequent imaging field of view (fov). Progress of dye uptake was monitored by snapshots of multiphoton images and pressure was adjusted to minimize swelling of the tissue.

Odor application started >1 h after dye injection. Food extract was prepared as described ^96^. Other odors (Sigma Aldrich) were prepared as 1000x stock solutions in deionized water (Fluka), vortexed, sonicated, and stored at −20°C. Fresh aliquots were diluted in ACSF to the final concentration before each experiment. For characterization of odor responses in interneurons (**Fig. 2; Fig. S2**), we used the following odor set (concentrations in µM): 10 Amino acid mix (equal parts of:), 10 L-Arginine (Arg), 10 eL-Lysine (Lys), 10 L-Phenylalanine (Phe), 10 L-Tryptophan (Trp), 10 L-Methionine (Met), 10 L-Histidine (His), 1 bile acid mix (Bmx; equal parts of: Taurodeoxycholic acid (TDCA), Taurocholic acid (TCA), Glycocholic acid (GCA)), 1 TDCA, 1 GCA, 10 nucleotide mix (Nmx; equal parts: Adenosine 5’-triphosphate (ATP) and Inosine 5’-monophosphate), 10 ATP. In each fov, each odor was applied twice in two separate sequences such that all odors were presented once before the second sequence started. The order of odors was newly randomized for each sequence and fov. For characterization of odor responses upon PIN (**Figs. 3, 7, 8; Fig. S3**), we used the following odor set (concentrations in µM): 10 Phe (F), 10 Trp (W), 10 Met (M), 10 Lys (K), 20 Phe + Met mix (F+M; equal parts), 20 Trp + Lys mix (W+K: equal parts), 1 bile acid mix (Bmx; equal parts of: TDCA, TCA, GCA), and 1:1000 dilution of food odor (Fex). In each fov, each odor was applied four times in four separate sequences. For each odor, two control and two PIN trials were interleaved in these four sequences. Two sequences were always paired such that a given odor was presented under 594 nm illumination (PIN) in the first sequence and under control conditions (Ctrl) in the second sequence or vice versa. The order of odors and photostimulation conditions was newly pseudo-randomized for the first sequence of each pair in each fov. Inter-stimulus intervals were between 2 and 3 min.

Odors were applied to the nasal epithelium for ∼3 s through a constant stream of ACSF using a computer-controlled, pneumatically actuated HPLC injection valve (Rheodyne, Rohnert Park, CA, USA) as described ^55^. One to four fovs were recorded in each fish (N = 12 fovs from five 212C-ArchTGFP fish; N = 19 fovs from eight dlx-eNpHR3.0YFP fish). Electrical stimulation of the mOT in conjunction with Ca^2+^ imaging (**Fig. 7**) was performed as described above, except that we applied ten stimuli (0.5 ms) at 5 Hz starting 0.2 s after the onset of 594 nm illumination in PIN trials. In each fov, a total of six stimulation amplitudes (5 V, 10 V, 20 V, 30 V, 50 V, and 75 V) were applied four times each. Ctrl and PIN trials were pseudo-randomized and paired as described above for odor stimulation. All these electrical stimulation experiments were performed following a bilateral bulbectomy. One to maximally six fovs were recorded in each fish (N = 12 fovs from five 212C-ArchTGFP fish; N = 18 fovs from six dlx-eNpHR3.0YFP fish).

### Image acquisition and optical stimulation (PIN)

In most experiments, multiphoton Ca^2+^ imaging in pDp was performed using a custom-built multiphoton microscope with a Galvo-Galvo scan head ^55^, a 20x water-immersion objective (NA 1.0, Zeiss), GaAsP photomultiplier tubes (PMT; Hamamatsu), and Scanimage/Ephus software ^94,97^. In each trial, images with 256 lines (256 pixels/line for rhod-2 imaging, and 512 pixels / line for OGB-1 imaging) were acquired at 128 ms per frame. After each trial, the field of view was readjusted to compensate for potential drifts using an automated routine that acquired a small z stack of ± 3 μm (step size, 0.5 μm). For rhod-2 imaging, fluorescence was excited at 860 nm. Red (rhod-2) and green (GFP) emitted light were detected simultaneously through bandpass filters (645/75 nm and 515/30 nm, respectively). For OGB-1 imaging, fluorescence was excited at 928 nm and emission was detected through a bandpass filter (535/50 nm) by a gated GaAsP PMT (Hamamatsu) that was further protected by a narrow blocking filter centered on 594 nm. The intensity of the 2-photon excitation light was adjusted in each fov to minimize photobleaching.

Optogenetic stimulation (PIN) during Ca^2+^ imaging was performed as described (Frank et al., 2019). Briefly, orange laser light (594 nm) was directed at pDp through an optical fiber (200 μm diameter; ThorLabs) positioned approximately 100 – 200 μm from the brain surface. Brief pulses of light (450 μs) were coupled into the fiber using a digital micromirror device (DMD; Texas Instruments; ^55^) and synchronized to every second line of image acquisition. Simultaneously, the PMT gate was switched off. After data acquisition, blanked lines were removed from images. The same processing step was applied to trials without photostimulation, resulting in final images with 128 lines and a fill fraction of approximately 40% under all conditions. The intensity of orange light at the tip of the fiber, averaged over the duty cycle, was 6 – 8 mW.

Most odor responses were measured in the core section of Dp between ∼100 µm and 260 µm from the ventral-most aspect (**Figs. 3, 7, 8; Fig. S3**). This section is densely innervated by processes of interneurons but contains few interneuron somata. To characterize odor responses in interneurons (**Fig. 2; Fig. S2**), images were acquired ∼50 – 85 µm or 50 – 120 µm from the most ventral aspect of Dp in 212C-GFP and dlx:GFP, respectively. At these depth, interneuron somata are more abundant than in the core section of Dp albeit still sparse compared to other neuron (unpublished observations). Subtle differences between odor responses measured in 212C-GFP and dlx:GFP fish (**Fig. 2; Fig. S2**) may be due to slight differences in imaging depth.

For co-expression analysis of 212C-mCherry and dlx:GFP, some images were acquired using a Resonant-Galvo scan head, excitation at 800 nm, and detection through bandpass emission filters (GFP: 515/30 nm; mCherry: 641/75 nm). Images were collected throughout the full dorso-ventral extent of pDp (∼300 µm).

### Analysis of Ca^2+^ signals

Images were registered across trials and regions of interest (ROIs) were drawn manually over all visible somata in each image fov using custom software (https://github.com/i-namekawa/Pymagor). For each ROI, we calculated the relative change in fluorescence ΔF/F. The baseline fluorescence F was averaged over a 1 s time prior to stimulus onset. ΔF/F traces were transformed into firing rate estimates by a deep learning-based spike inference algorithm (“Cascade”. ^60^) using the model “OGB_zf_pDp_7.5Hz_smoothing200ms_causalkernel” (available at https://github.com/HelmchenLabSoftware/Cascade) for both rhod-2 and OGB-1 data sets.

To quantify responses of neurons to odors or electrical stimulation (neuron-odor or neuron-stimulus pairs), inferred firing rates were averaged over a temporal response window of 3 s in odor stimulation experiments, starting with the onset odor stimulation, unless stated otherwise. In electrical stimulation experiments, a longer response window of 11 s was chosen to include the prolonged activity in response to high stimulus amplitudes (**Fig. 7**). Neurons that showed no response to at least one odor or electrical stimulus were excluded from further analyses; we used a threshold of 0.128 for the summed number of estimated spikes as a - conservative - criterion.

### Network model

#### Model

The simulation of pDp consists of 4000 E neurons, 500 FFI neurons and 500 FBI neurons that were modeled as leaky integrate-and-fire units with conductance-based synapses. A spike emitted by neuron *y* from population Y triggers an increase in the conductance *g_yx_* in the postsynaptic neuron,*x*:

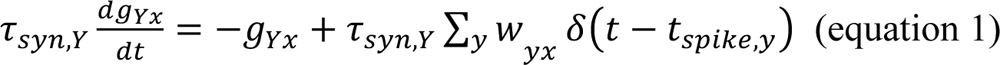

Conductance changes triggered by the OB and local populations P affect the membrane potential of neuron *x* which evolves according to equation (2).

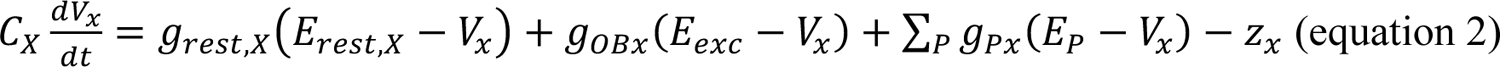

When the membrane potential reaches a threshold V_th_, the neuron emits a spike and its membrane potential is reset and clamped to *E_rest_*) during a refractory period τ_ref_.

Excitatory neurons are endowed with adaptation with the following dynamics ^98^

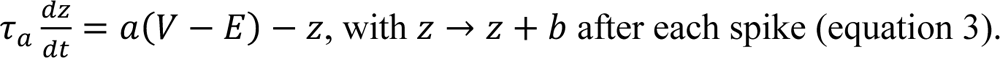

Neuronal parameters were similar to ^18^ (Table 1). The time constants of inhibitory and excitatory synapses (τ*_syn,l_* and τ*_syn,E_*) were 10 ms and 30 ms, respectively.

To show that the results were consistent across a wide range of parameters, we simulated 10 sparsely connected random networks with different connection probabilities p_YX_ and synaptic strengths w_YX_ as summarized in Table 2. Connections between neurons were drawn from a Bernoulli distribution with probability p_YX_. Synaptic strengths w_YX_ were then fitted to reproduce observations in *ex-vivo* pDp, as described in ^18^. Simulations were performed using Matlab. Differential equations were solved using forward Euler and an integration time step of dt = 0.1 ms.

**Table 1.**
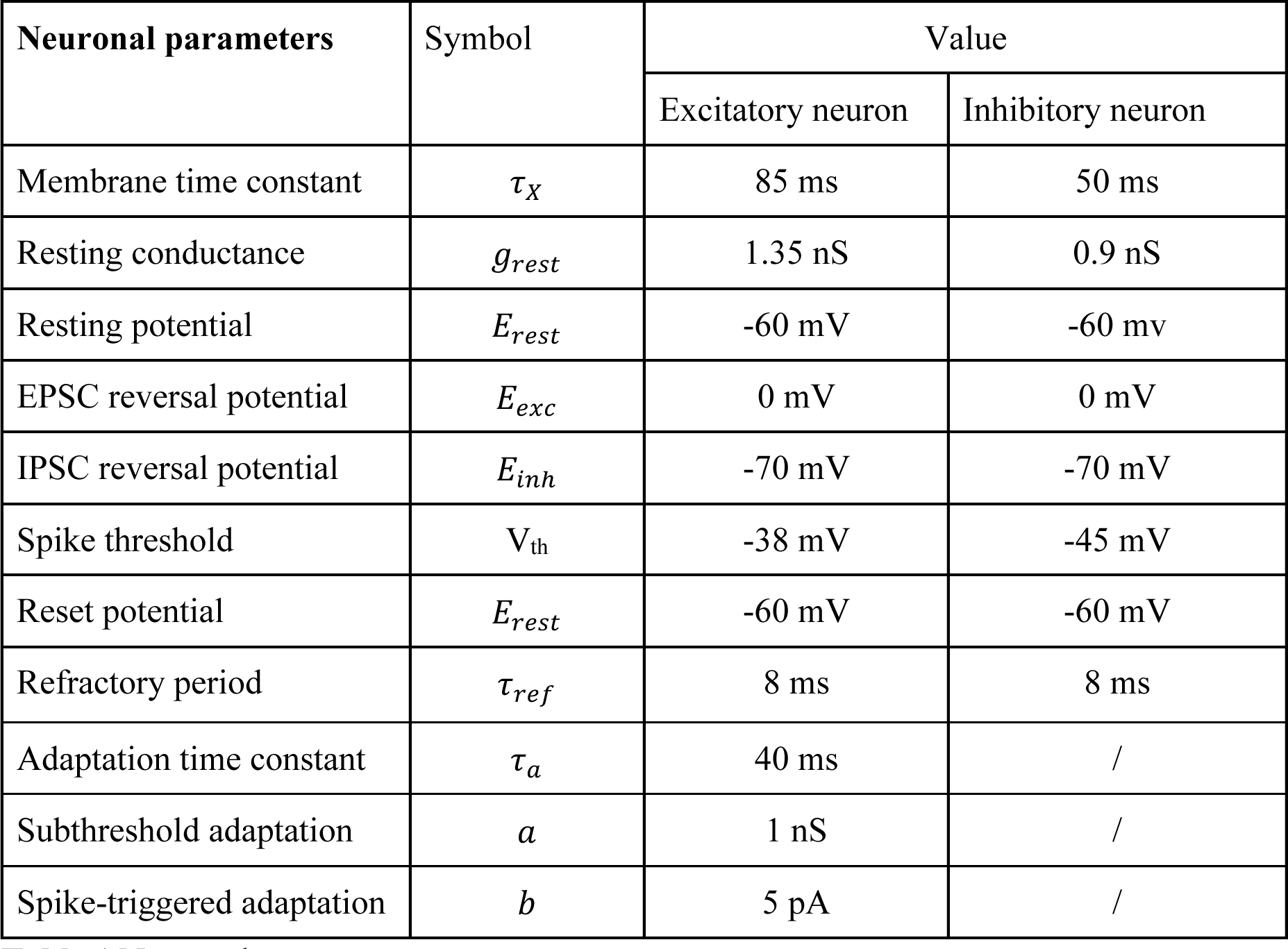
Neuronal parameters.

**Table 2.**
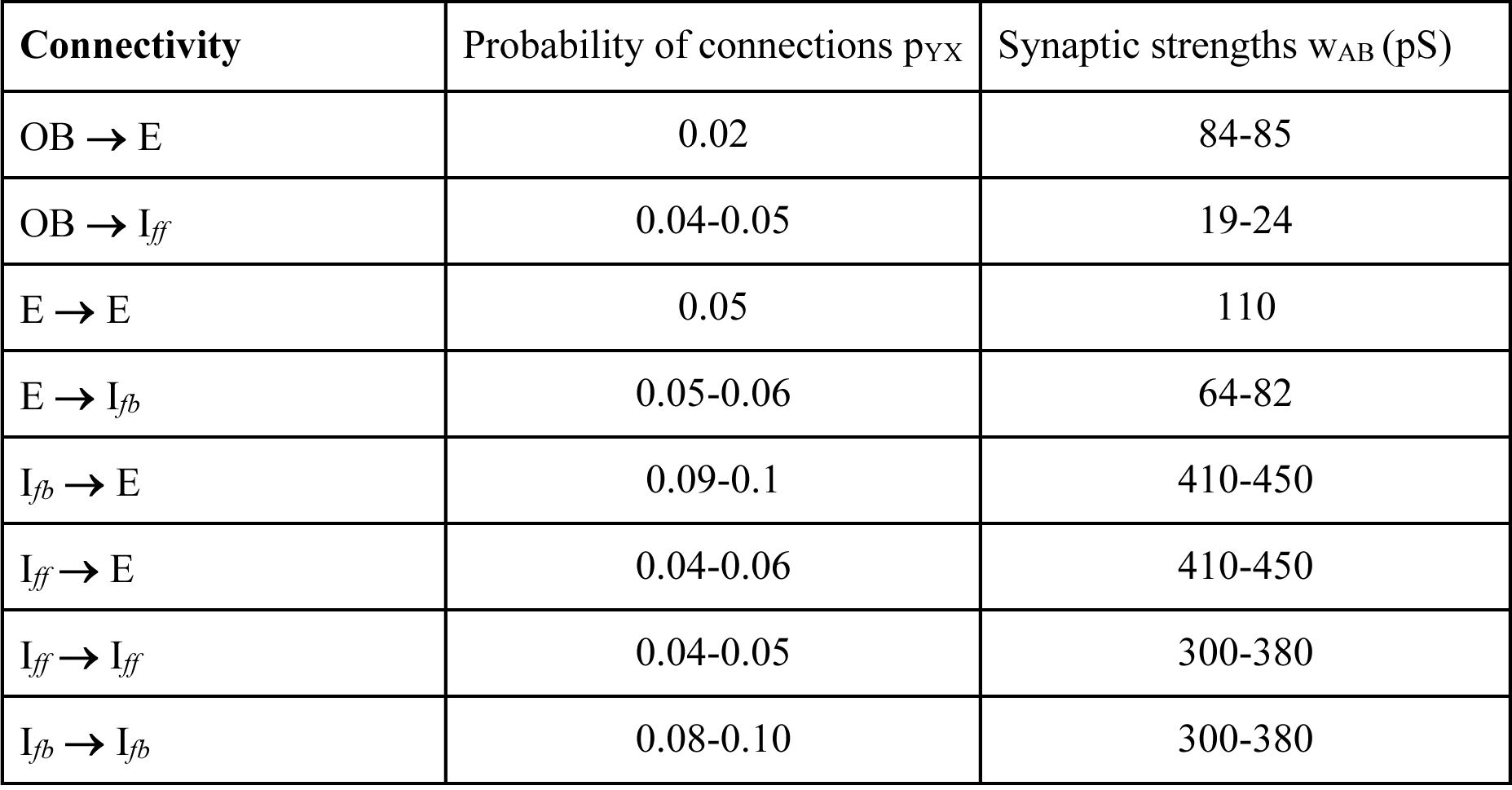
Values of the connectivity parameters used (for simplicity, wIff*-*Iff= wIfb*-*Ifb and wIff*-*E= wIfb*-*E)

| Connectivity | Probability of connections $p_{yx}$ | Synaptic strengths $w_{AB}$ (pS) |
| --- | --- | --- |
| OB → E | 0.02 | 84-85 |
| OB → $I_{ff}$ | 0.04-0.05 | 19-24 |
| E → E | 0.05 | 110 |
| E → $I_{fb}$ | 0.05-0.06 | 64-82 |
| $I_{fb} \rightarrow$ E | 0.09-0.1 | 410-450 |
| $I_{ff} \rightarrow$ E | 0.04-0.06 | 410-450 |
| $I_{ff} \rightarrow I_{ff}$ | 0.04-0.05 | 300-380 |
| $I_{fb} \rightarrow I_{fb}$ | 0.08-0.10 | 300-380 |

#### Afferent input

Each E and FFI neuron received external excitatory input from a pool of 1500 mitral cells. During baseline activity, mitral cells fired at 6 Hz. During odor presentation (2 s) firing rates of 75 mitral cells were decreased (“inhibited”): rates were drawn from a discrete uniform distribution between 0 to 5 Hz, and the onset latency was drawn from a discrete uniform distribution between 0 to 200 ms. At odor onset, firing rates of 150 mitral cells increased (“activated” cells). Their firing rates were drawn from a uniform distribution ranging from 8 to 32 Hz and the onset latency was drawn from a uniform distribution between 0 to 200 ms. After odor onset, their firing rate decreased back to baseline with a time constant of 1, 2 or 4 s (equally distributed). Spike trains were then generated by Poisson processes.

#### Assemblies

Unless stated otherwise we created 20 assemblies (“odor memories”) in each network. For each learned odor, we selected the 60 to 100 E neurons (fixed number for each network) that received the highest number of connections from activated mitral cells. We then created additional connections between these assembly E neurons and eliminated existing connections between other neurons and assembly E neurons to maintain a constant number of E input connections per E neuron. The connection probability between assembly E neurons was therefore in average 5-fold higher than the probability among E neurons outside assemblies. We then selected the 10 – 25 FBI neurons (fixed number for each network) that received the highest number of connections from assembly E neurons and increased the probability of connection from these assembly I neurons to assembly E neurons by a factor 10 in average, resulting in an EI assembly containing both E and I neurons. Similar to E connections, we eliminated existing connections between non-assembly I neurons and assembly E neurons to maintain a constant number of I input connections per E neuron. In total, 5 sets of 2 *rand* and 4 *struct* networks which shared the same OBèE, OBèI, IèI and EèI connectivity were simulated (2 *struct* networks per rand network, as 2 different sets of 20 learned odors were used to create assemblies).

Odors presented to simulated networks were generated by random selection of mitral cells as described and did not match assemblies. This procedure mimicked an experimental setting in which an animal with an unknown history is presented random odor stimuli.

#### Inhibiting the inhibitory neurons

To mimic PIN, we deleted the output connections of a random subset of inhibitory neurons 500 ms before odor onset until the end of the odor presentation. 34% of the FFI or 51% of the FBI were inhibited. The percentage of inhibited neurons was set to obtain an average modulation (firing rate during vPIN normalized by control firing rate) as close as the one observed experimentally by PIN_212C_ and PIN_dlx_ in pDp.

#### Analysis

Unless stated otherwise, odor responses were time-averaged over the first 1.5 seconds of odor presentations, averaged over 20 different odors, and then averaged over neurons.

### Data analyses (experiments and simulations)

#### Lifetime and population sparseness

Lifetime and population sparseness were calculated using the metric ^99^:

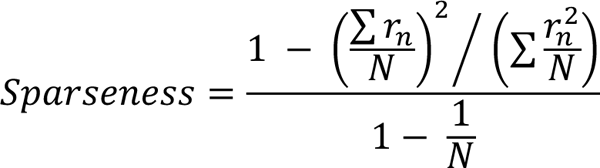

This normalized metric describes the ‘peakiness’ of a distribution and ranges between 0 (all responses equal) and 1 (only one non-zero response). Lifetime sparseness: distribution of response amplitudes across odors in single neurons. Population sparseness: distribution of response amplitudes to a single odor across neurons. (Vinje and Gallant, 2000)

#### Gain index

The gain index was defined as the ratio between the mean firing rates of E neurons during vPIN or PIN and under control conditions. This index is not a direct measure of total inhibition because inhibition was only partially eliminated during vPIN/PIN. Nonetheless, it provides an experimentally accessible measure of the “inhibitory gain” that counteracts recurrent amplification in the balanced state. Normalization by the mean firing rate under control conditions avoids problems with division by firing rates which can be close to zero.

#### Linear regression

Linear fits (**Figs. 3, 4, 7, 8**) were performed using a total least squares procedure, which minimizes the orthogonal distance of the regression line from the data points. Thus, this method does not distinguish between dependent and independent variables and accounts for the equality of measurement errors in the two variables.

#### Contribution to *r*

The contribution of individual neurons to the correlation between activity patterns is determined by the corresponding element in the Pearson correlation, which is the sum over the individual contributions of all neurons. For each simulated network, the odor pairs for which pattern correlations increased strongly upon vPIN_FBI_ (Δ*r* = *r*_PIN_ - *r*_Ctrl_ > 0.25) were selected and for each pair of odor-response pattern, 2% of neurons (80) were randomly selected. In **Fig 8**, all odor pairs are shown.

### Statistics

Unless stated otherwise, sample means are reported ± their standard error (± s.e.m.). N indicates the number of fovs, and, unless stated otherwise, n indicates number of neurons. No statistical methods were used to predetermine sample sizes, but our sample sizes are similar to those in prior reports and are typical for the field. To test whether the linear correlation between two variables was statistically significant, we performed a t test of the null hypothesis that the observed correlation coefficient r comes from a population with r = 0. To test whether the regression slope of linear fits was significantly larger than 1 (**Fig. 3**), we used bootstrapping to estimate the sampling distribution of the regression slope (sampling with replacement, repeated 10’000 times). All statistical tests were two-sided unless stated otherwise, and p < 0.05 was considered statistically significant. Significance levels are indicated as follows: p ≥ 0.05: ns; p < 0.05: *; p < 0.01: **; p < 0.001: ***.

## DATA AND CODE AVAILABILITY.

Matlab code to simulate pDp_sim_ is available at https://github.com/clairemb90/pDp-model.

## ACKNOWLEDGEMENTS

We thank Tetsuya Koide, Nobuhiko Miyasaka, and Yoshihiro Yoshihara for sharing data on the SAGFF(LF)212C line, which is available through the National BioResource Project “Zebrafish” in Japan. We thank Andreas Lüthi for critical comments on the manuscript and members of the Friedrich group for insightful discussions. We thank Enrico Tagliavini for IT support. This work was funded by the Novartis Research Foundation, by the European Research Council (ERC) under the European Union’s Horizon 2020 research and innovation program (grant agreement no. 742576), by the Swiss National Science Foundation (grants no. 31003A_172925/1, 310030B_152833/1, PCEFP3_202981), the Max Planck Society, the German Research Foundation (DFG) under Germany’s Excellence Strategy (EXC 2067/1-390729940), the Human Frontiers Science Program (HFSP; fellowship to T.F.), and the European Molecular Biology Organization (EMBO; fellowship to T.F.).

## AUTHOR CONTRIBUTIONS

Conceptualization: RWF, TF and CMB. Experimental design: RWF and TF. Resources (transgenic lines): TF and EAB. Experiments: TF and BJ. Simulations: CMB (lead) and FZ (supporting). Data analysis: TF, CMB, PR and RWF. Writing: RWF, TF and CMB. Review and Editing: all authors.

## SUPPLEMENTAL INFORMATION

**Figure S1.**
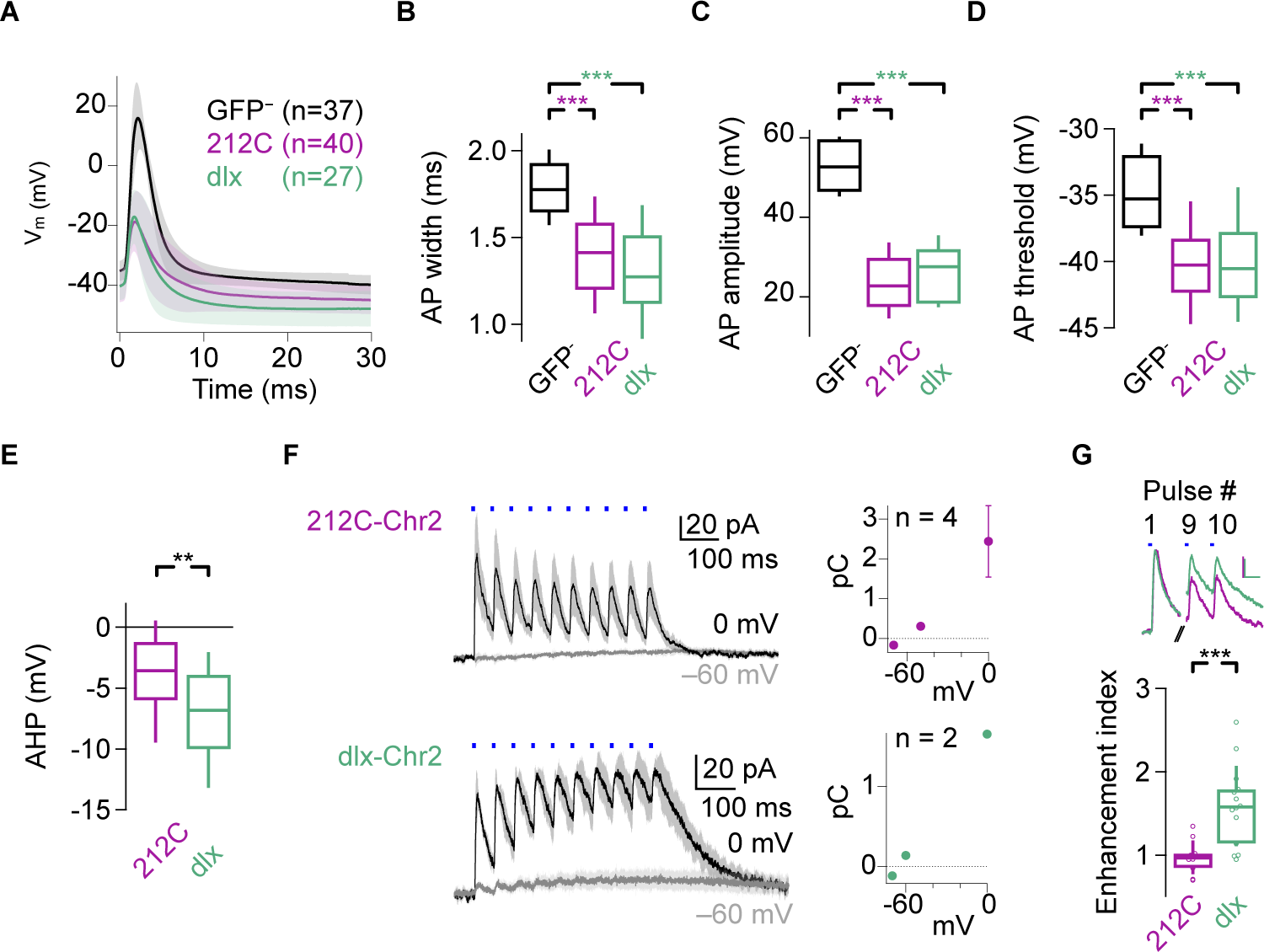
Further characterization of two populations of inhibitory interneurons in Dp. **(A)** Mean action potential waveforms (± s.d.) recorded from 212C-GFP^+^, dlx:GFP^+^, and GFP^−^ neurons (n: number of neurons). **(B)** Comparison of action potential width; action potential width was measured as the time between the two zero crossings of the second derivative of the membrane voltage, before and after the action potential peak, respectively (Kruskal-Wallis test, n = 102, p < 10^−9^). Nonparametric comparisons with GFP^−^ (n = 37): 212C-GFP^+^, p < 10^−6^, n = 39; dlx:GFP^+^, p < 10^−7^, n = 26). **(C)** Action potential amplitude (Kruskal-Wallis test, n = 102, p < 10^−15^). Nonparametric comparisons with GFP^−^ (n = 37): 212C-GFP^+^, Q = 7.56, p < 10^−13^, n = 39; dlx:GFP^+^, Q = 6.11, p < 10^−8^, n = 26). **(D)** Action potential threshold (Kruskal-Wallis test, n = 103, p < 10^−9^). Nonparametric comparisons with GFP^−^ (n = 37): 212C-GFP^+^, p < 10^−7^, n = 39; dlx:GFP^+^, p < 10^−5^, n = 27). **(E)** Action potential afterhyperpolarization (AHP) amplitude (Wilcoxon rank-sum test: p = 0.009; 212C-GFP^+^, n = 42; dlx:GFP^+^, n = 27). **(F)** Left: Mean (± s.e.m.) IPSCs and EPSCs in GFP^−^ neurons in response to trains (10 pulses at 20 Hz) of 0.5 ms full-field blue light stimulation (blue bars) in 212C-Chr2YFP (top) and dlx-Chr2YFP fish (bottom; same data as in Fig. 1D but showing complete trains). Right: Average (± s.e.m.) charge transfer over the first 50 ms (first “pulse”) as a function of holding potential (n: number of neurons). **(G)** Top: IPSCs number 1, 9 and 10 evoked by trains of blue light pulses in GFP^−^ neurons (same as Fig. 1E). Bottom: charge transfer evoked by the last two pulses, normalized to the first pulse (Wilcoxon rank-sum test: p = 0.0009; 212C-GFP^+^, n = 10; dlx:GFP^+^, n = 14). In dlx-Chr2YFP fish, charge transfer increased because IPSCs broadened.

**Figure S2.**
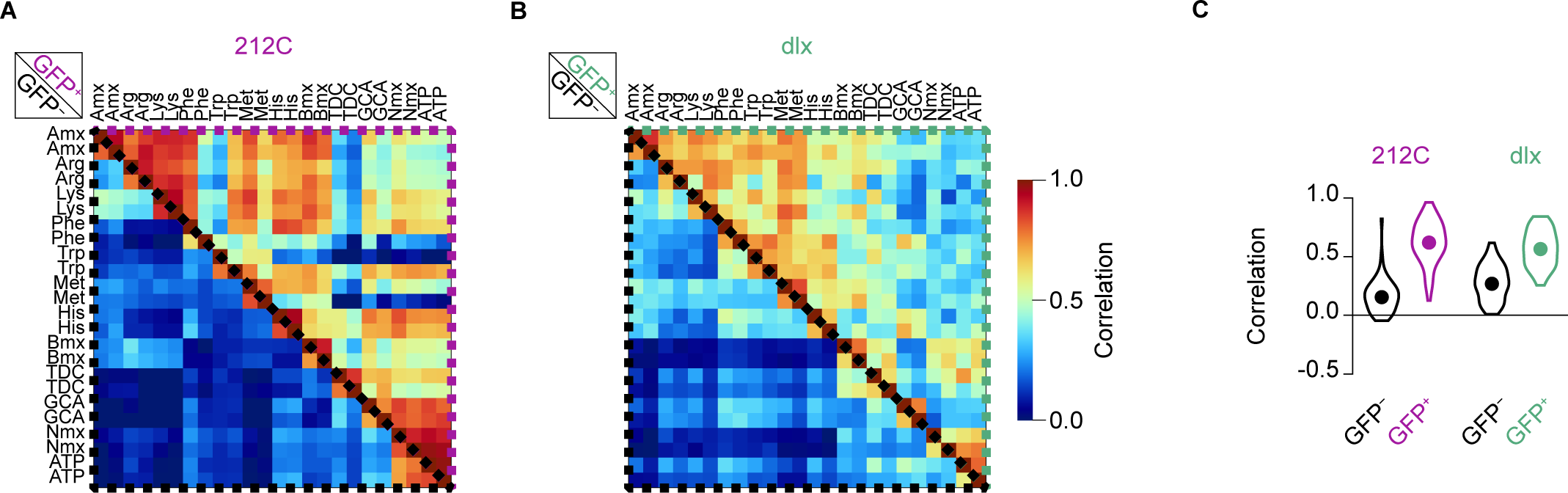
Population analysis of odor-evoked responses in populations of putative principal cells and interneurons in pDp. **(A)** Pearson correlation between odor-evoked activity patterns in simultaneously recorded putative principal cells (GFP^−^ neurons, below diagonal) and interneurons (GFP^+^ neurons, above diagonal) in 212C-GFP fish (n = 1515 GFP^−^ and n = 50 GFP^+^ neurons from N = 5 fovs; two trials per odor). **(B)** Same for dlx:GFP fish (n = 1750 GFP^−^ and n = 65 GFP^+^ neurons from N = 12 fovs). **(C)** Distribution of Pearson correlations between all pairwise activity patterns in. Two trials with each odor were averaged (Wilcoxon signed rank test; 212C-GFP fish: n = 66 odor pairs, p < 10^−19^; dlx:GFP fish: n = 66 odor pairs, p < 10^−16^).

**Figure S3.**
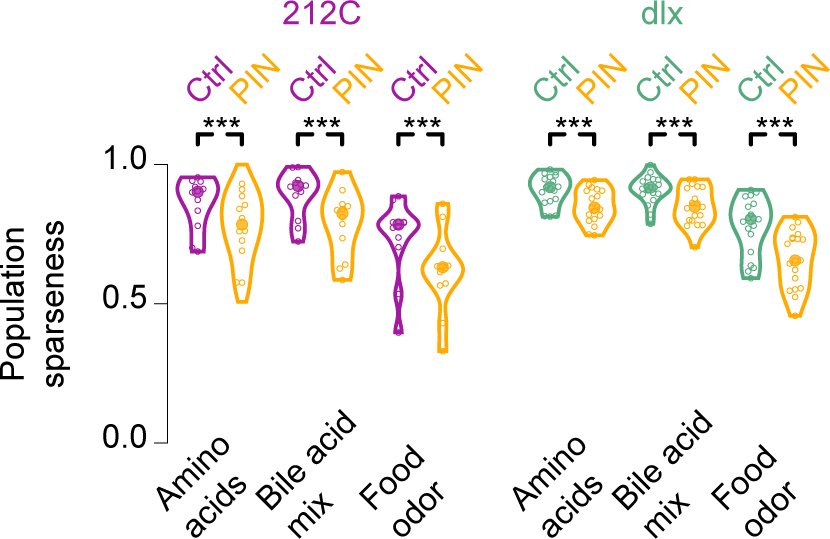
Effects of PIN on population sparseness of odor responses in pDp. Population sparseness of responses evoked by different classes of odors (amino acids: four individual amino acids and two binary mixtures; bile acids: one 3-component mixtures; one food extract; see Methods for details) under control conditions (Ctrl) and during vPIN_212C_ (left) and vPIN_dlx_ (right). Datapoints represent individual fovs (Wilcoxon signed rank tests: 212C: N = 12 fovs, p = 0.0005 for all three cases: dlx: N = 19 fovs; amino acids: p < 10^−4^; bile acid mix: p = 0.0002; food extract: p < 10^−4^).

**Figure S4.**
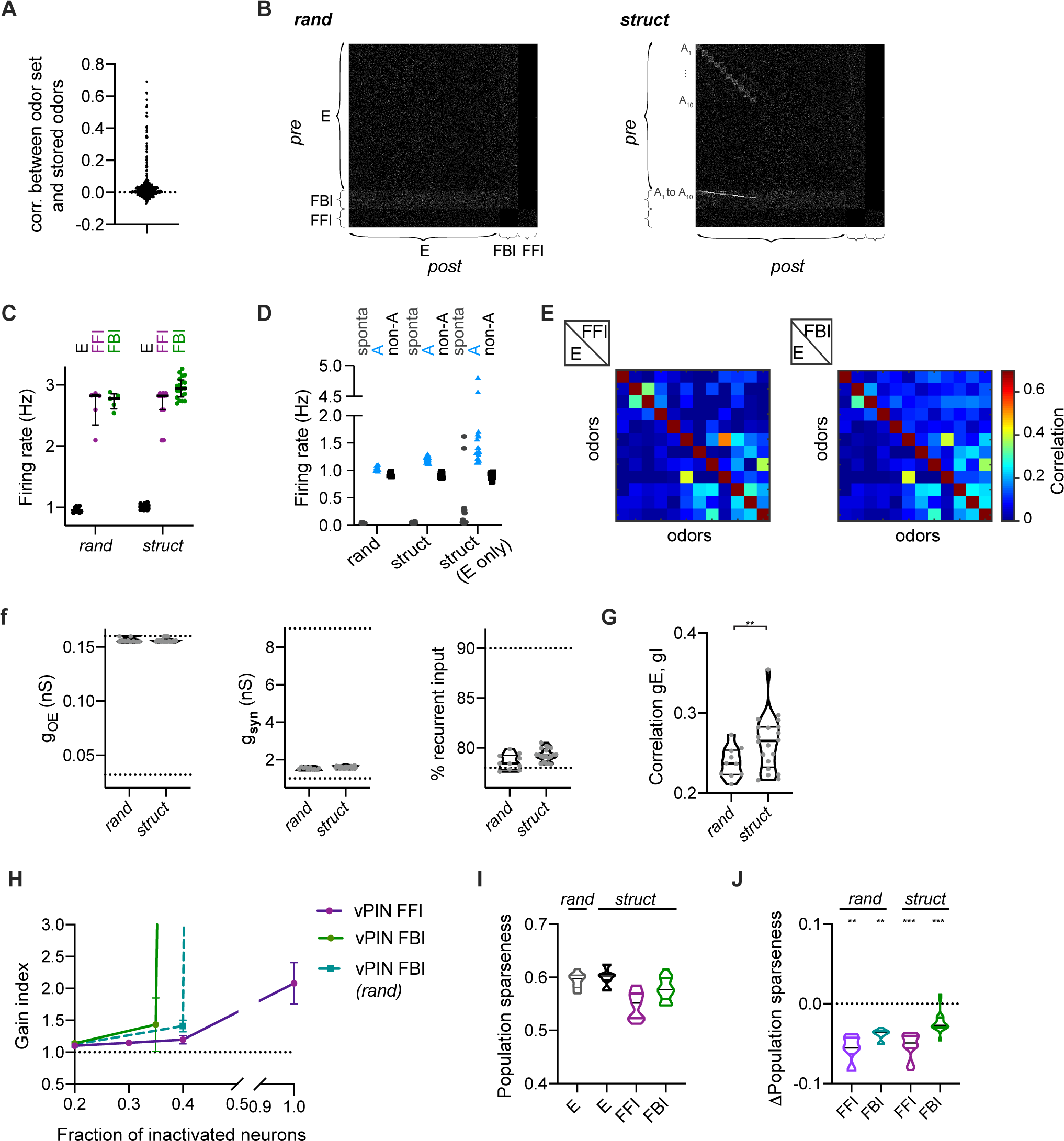
Computational model of pDp: additional results. **(A)** Pearson correlation between afferent patterns presented to pDp_sim_ (n=20) and learned afferent patterns used to create EI assemblies (n = 20). **(B)** Example connectivity of a *rand* network and a corresponding *struct* network. A white dot indicates the presence of a connection between neurons. Half of the network is depicted, including 10 out of 20 assemblies (A), which consists of 80 E and 10 FBI neurons each. **(C)** Mean firing rates of E, FFI and FBI neurons in *rand* and *struc*t networks during presentation of an odor. **(D)** Firing rates averaged during spontaneous (sponta) activity or during odor presentation, over neurons belonging to an assembly (A) and the remaining ones (non-A). **(E)** Correlations between activity patterns evoked by 12 odors in E (lower triangles) and FFI or FBI (upper triangles). Example of 1 network. **(F)** Left: averaged odor-evoked afferent conductance. Middle: odor-evoked synaptic conductance. Right: percentage of E input coming from recurrent connections during odor presentation. The experimental range measured in *ex-vivo* Dp is delineated by the dotted lines. **(G)** Co-tuning, quantified by the correlation between time-averaged E and I conductances in each neuron in response to various odors (average across neurons, n = 10 and 20 *rand* and *struct* networks, respectively; Wilcoxon matched-pairs signed rank test: p = 0.002). **(H)** Gain index as a function of the fraction of inactivated neurons. **(I)** Population sparseness of responses evoked by different odors. **(J)** Changes in population sparseness induced by vPIN (vPIN-Ctrl, one sample Wilcoxon signed rank test: FFI, rand: p = 0.002; FBI, rand: p < 0.002; FFI, struct: p < 0.0001; FBI, struct: p < 0.0001).

**Figure S5.**
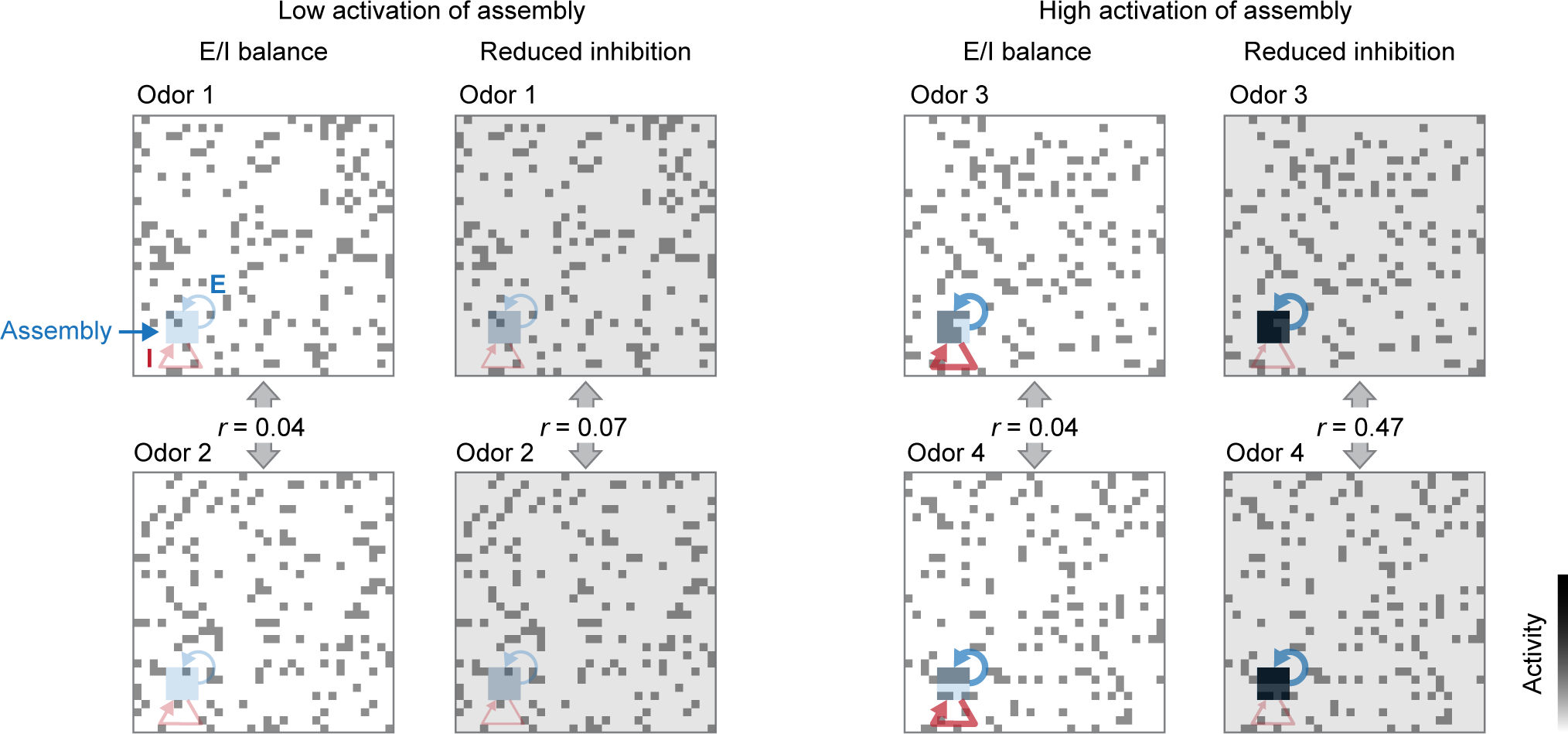
Mechanism generating runaway correlations: schematic illustration. Grids represent 32 x 32 E neurons; shaded square depicts an EI assembly; arrows represent feedback excitation (E) and multisynaptic feedback inhibition (I). Left: odors 1 and 2 are uncorrelated and activate a small subset of neurons within the assembly. Reducing inhibition enhances activity within the assembly slightly more than outside the assembly but the resulting increase in pattern correlation remains small. Right: odors 3 and 4 are also globally uncorrelated but activate a larger subset of neurons within the assembly. Because feedback gain increases with assembly activation (non-linear amplification), a reduction in inhibition strongly enhances activity within the assembly. As a consequence, the global pattern correlation becomes high even though activity outside the assembly is uncorrelated. This “runaway correlation” does not occur when excitation and inhibition are precisely balanced because nonlinear amplification within assemblies is canceled. In poorly balanced networks, runaway correlations therefore emerge in response to subsets of inputs (odors) depending on the precise relation between input patterns (odors) and pre-existing memories (assemblies). Note that this is a schematic illustration with fewer neurons and assemblies than the biologically constrained simulation.

